# Functional characterization of eQTLs and asthma risk loci with scATAC-seq across immune cell types and contexts

**DOI:** 10.1101/2023.12.24.573260

**Authors:** Julong Wei, Justyna Resztak, Ali Ranjbaran, Adnan Alazizi, Henriette E Mair-Meijers, Richard Slatcher, Samuele Zilioli, Xiaoquan Wen, Francesca Luca, Roger Pique-Regi

## Abstract

Cis-regulatory elements (CREs) control gene transcription dynamics across cell types and in response to the environment. In asthma, multiple immune cell types play an important role in the inflammatory process. Genetic variants in CREs can also affect gene expression response dynamics and contribute to asthma risk. However, the regulatory mechanisms underlying control of transcriptional dynamics across different environmental contexts and cell-types at single cell resolution remains to be elucidated. To resolve this question, we performed scATAC-seq in activated peripheral blood mononuclear cells (PBMC) from 16 children with asthma with phytohemagglutinin (PHA) or lipopolysaccharide (LPS), and treated with dexamethasone (DEX), an antiinflammatory glucocorticoid. We analyzed changes in chromatin accessibility, measured transcription factor motif activity, and identified treatment and cell-type specific transcription factors that drive changes in both gene expression mean and variability. We observed strong positive linear dependence between motif response and their target gene expression changes, but negative in variability changes. This result suggests that an increase of transcription factor binding tightens the variability of gene expression around the mean. We then annotated genetic variants in chromatin accessibility peaks and response motifs followed by computational fine-mapping of eQTL signals from a pediatric asthma cohort. We found that eQTLs were 5-fold enriched in peaks with response motifs and refined the credible set for 410 asthma risk genes, with 191 having the causal variant in response motifs. In conclusion, scATAC-seq enhances the understanding of molecular mechanisms for asthma risk variants mediated by gene expression.

## Introduction

Glucocorticoids are anti-inflammatory drugs widely used to treat asthma (Jenkins et al. (2020)), autoimmune diseases (Snider Potter (2011); Mithoowani Arnold (2019); Hughes et al. (2017); Mieli-Vergani et al. (2018); Wang et al. (2018); Strum et al. (2020)) or other inflammatory conditions (Bruscoli et al. (2022)). Glucocorticoids main mechanism of action is through binding and activation of the glucocorticoid receptor (GR) which then translocates to the nucleus and activates the anti-inflammatory response through two main mechanisms: 1) the activated receptor binds glucocorticoid response elements (GREs) thus activating the expression of anti-inflammatory genes; 2) the activated receptor inhibits pro-inflammatory transcription factors (TFs), e.g NFKB and AP-1, from binding their target genes thus repressing the expression of inflammatory genes (Silverman et al. (2005)). We previously performed scRNA-seq in peripheral blood mononuclear cells (PBMCs) of children with asthma to systematically decipher transcriptome dynamic changes in response to glucocorticoids in a cell type specific way (Resztak et al. (2023)). However, the effect of glucocorticoids on the chromatin conformation landscape in different immune cell types has not been characterized yet at single cell resolution, and we don’t have a full understanding of which TFs are involved in modulating the immune response in combination with glucocorticoids.

Genome-wide association studies (GWAS) have identified thousands of genetic variants associated with human immune-mediated diseases (Claussnitzer et al. (2020); Tsuo et al. (2022)). The majority of these disease-associated loci reside in noncoding regions of the genome (Maurano et al. (2012)). These noncoding genetic variants contribute to disease through effects on gene regulation which ultimately lead to gene expression changes (Nicolae et al. (2010)). Transcriptome-wide association studies have enabled significant progress in associating gene expression changes to disease risk through genetic effects on gene expression (Gamazon et al. (2015)); yet there is still a substantial gap between disease-associated variants and eQTL signals, with more than half of GWAS signals not colocalized with any eQTLs in GTEx (Aguet et al. (2020)). It has been suggested that only a small proportion of disease heritability (about 11% across traits) is mediated by the cis-genetic component of gene expression (Yao et al. (2020)). A potential explanation is that eQTLs that are important for disease risk only exhibit their effects in context-specific ways, such as in specific cell types or environmental exposures (Findley et al. (2021, 2019); Moyerbrailean et al. (2016b); Resztak et al. (2023); Randolph et al. (2021); Oelen et al. (2022); Aquino et al. (2023)).

Genetic variants can affect gene expression by altering the affinity of TFs binding to the cis-regulatory elements, such as enhancers and promoters, in a cell type specific or environmental specific way (Degner et al. (2012); Moyerbrailean et al. (2016a); Albert Kruglyak (2015); Nasser et al. (2021); Boix et al. (2021)), thus contributing to disease risk (Figure 1A). Previous studies have shown that eQTL signals from bulk tissue are enriched in cis-regulatory elements (Aguet et al. (2020)), especially in regions of accessible chromatin that are captured in the most abundant cell type (Zhang et al. (2021)). The various environmental factors, including immune stimuli or drug treatments, interact with genetic variants via activating specific TFs (Findley et al. (2019)). Previously, we demonstrated that ATAC-seq can be used to determine transcription factor binding motifs associated with specific treatments and to annotate genetic variants in these motifs in endothelial cells (Findley et al. (2019)). Integration of these environment-specific regulatory features with eQTL data can improve the discovery of genes with eQTLs (eGenes) and prioritize putatively causal variants, thus achieving a better understanding of the mechanism of context-specific genetic regulation of gene expression (Findley et al. (2019)).

**Figure 1.**
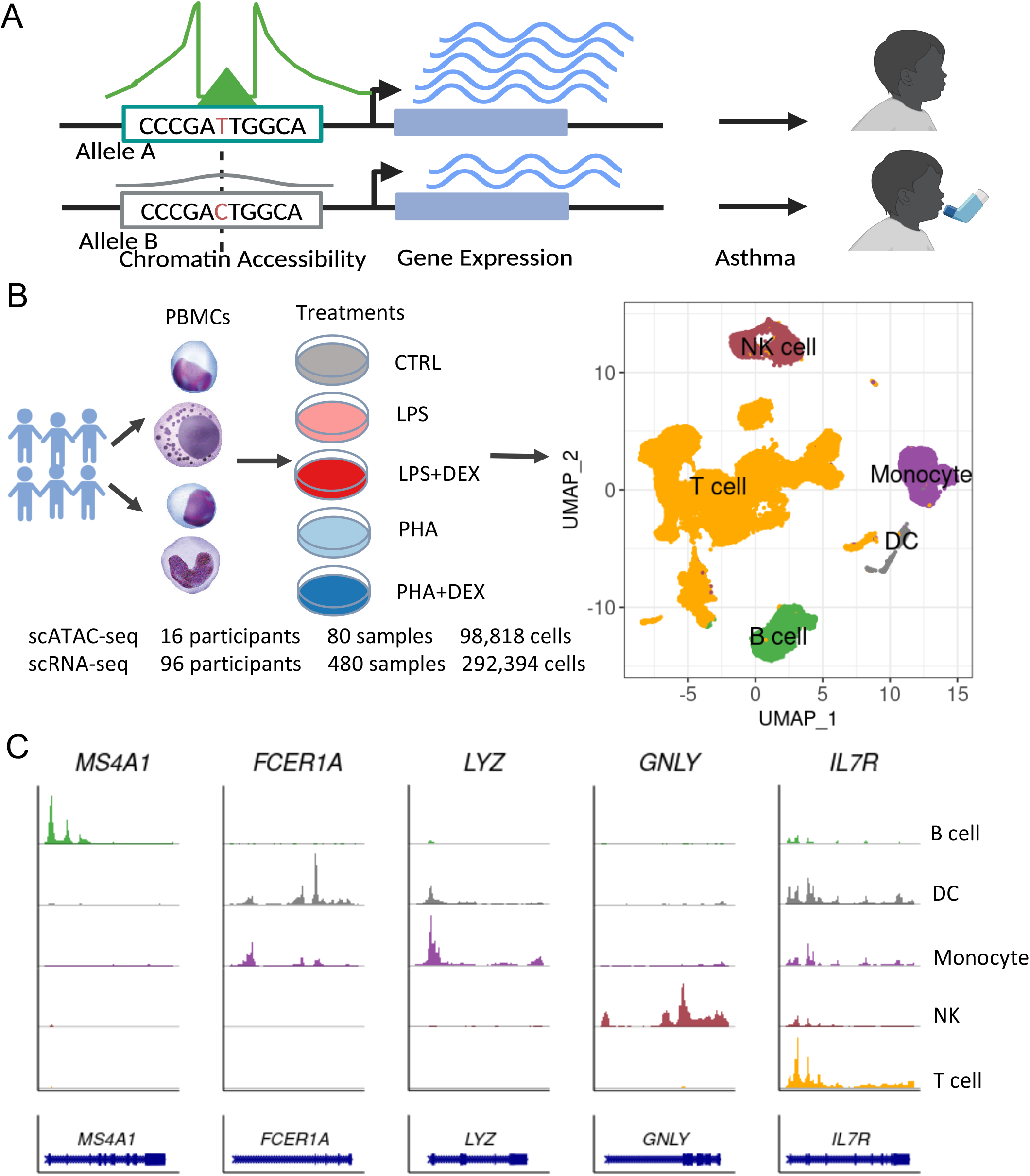
Study design. **A** Overview of the study connecting chromatin accessibility and gene expression from single cell experiments, and genetic variants associated with asthma. **B** Experimental data collected using scATAC-seq data for 16 participants stimulated with lipopolysaccharide (LPS), phytohemagglutinin (PHA) and treated with dexamethasone (DEX), and previously available scRNA-seq data. UMAP visualization of the 98,818 high quality cells colored by cell types, B cell (green), Monocyte (purple), NK cells (maroon), T cells (orange) and Dendritic cells (DCs, grey). **C** Normalized chromatin accessibility of the cis-regulatory elements (CRE) nearby the cell type-specific marker genes, *MS4A1* (B cells), *FCER1A* (DC cells), *LYX* (monocyte), *GNLY* (NK cell) and *IL7R* (T cell).

To better understand the regulatory mechanisms underlying genetic control of transcriptional dynamics in asthma (Figure 1A), we collected PBMCs samples from 16 African American children with asthma from the ALOFT cohort (Asthma in the Life Of Families Today). We activated the PBMCs with lipopolysaccharide (LPS, a component of the bacterial membrane) or phytohemagglutinin (PHA, a T cell mitogen) and treated with the glucocorticoid dexamethasone for a total of 5 conditions (including the unstimulated control, Figure 1B). PHA acts as a general T cell mitogen, which induces T cell activation and division by binding to TCR/CD3 complex (Gulden et al. (2023); Faguet (1977)). PHA binds multiple components of the TCR complex, leading to intracellular phosphorylation of ITAM motifs and activation of downstream signaling molecules (Schneider et al. (2012)). Consequently, PHA stimulates a diverse population of lymphocytes, potentially including all those relevant to asthma pathophysiology and not limited only to the “high Th2” asthma subtype (Hammad Lambrecht (2021)). This ensures a broad lymphocyte-mediated proinflammatory state that allows cellular communication between the innate and adaptive immune cells. LPS, a well-studied environmental antigen, is known to activate the acute inflammatory response by inducing TLR signaling (Ngkelo et al. (2012)). Previous studies have shown that during asthma exacerbation the circulating inflammatory cells demonstrate an enrichment of asthma-associated gene networks involved in myeloid cell activation and differentiation. LPS acts as a strong upstream driver of these signatures (Jones et al. (2023)). We have selected these immune stimulatory agents to capture a broad immune activation state and properly allow for the cross-talk between the adaptive and innate immune cells in peripheral blood mononuclear cells from asthma patients.

We performed scATAC-seq to study the chromatin accessibility changes and relevant transcription factor binding motifs across cell types and treatment conditions. We integrated the scATAC-seq data with previously collected scRNA-seq data from 96 ALOFT participants (Resztak et al. (2023)) to study the regulatory mechanisms underlying gene expression changes in mean and variability. Finally, we derived a regulatory annotation and used it to computationally fine-map eQTLs, which allowed us to further characterize asthma associated loci using colocalization and TWAS. We demonstrated that using a context-aware functional annotation increases our ability to fine-map and molecularly dissect risk loci for asthma.

## Material and Methods

### Biological samples and genotyping

Peripheral blood mononuclear cells (PBMCs) used in this study were collected as part of the longitudinal study Asthma in the Life of Families Today (ALOFT, recruited from November 2010 to July 2018, Wayne State University Institutional Review Board approval #0412110B3F) (Resztak et al. (2021)) from children 10-17 years old. For this study we focused on participants that self-reported as Black (this group experiences higher asthma severity and mortality and is generally understudied), and were 10-16 years old. The demographic information of individuals for scRNA-seq and scATAC-seq are summarized in the Tables (Table S1 and S2). PBMCs were extracted using a previously-published Ficoll centrifugation protocol (Weckle et al. (2015)), cryopreserved in freezing media and stored in liquid nitrogen until the day of the experiment. All individuals in this study were genotyped from low-coverage (∼ 0.4X) whole-genome sequencing and imputed to 37.5 million variants using the 1000 Genomes database by Gencove (New York, NY).

### Cell culture and single-cell experiments

Cell culture and treatment was performed following the protocol previously described by Resztak et al. (2023). Cells from 16 donors were processed in parallel in independent wells. Thawed PBMCs were incubated in starvation media overnight (90% RPMI 1640, 10% CS-FBS, 0.1% Gentamycin, approx. 16 hours) and subsequently treated with either: 1 *µ*g/ml LPS+ 1 *µ*M dexamethasone (LPS+DEX), 1 *µ*g/ml LPS + vehicle control (LPS), 2.5 *µ*g/ml PHA + 1 *µ*M dexamethasone (PHA+DEX), 2.5 *µ*g/ml PHA + vehicle control (PHA), or vehicle control alone (control). The vehicle control used was ethanol, which is the solvent for dexamethasone, at a final concentration of 1:10,000 ml of media.The treatment concentrations are the same as previously used (Barreiro et al. (2010); Moyerbrailean et al. (2016b)). After six hours, cells were pooled across individuals within each treatment condition, for a total of 5 experimental pools. Each pool was split in half to perform single cell RNA-seq and single cell ATAC-seq in parallel. The scRNA-seq data were previously published (Resztak et al. (2023)) and available at dbGAP accession number phs002182. Each scATAC-seq pool underwent nuclei isolation and was loaded on 2 channels of the 10x Genomics®Chromium Instrument, for a total of two loading batches of five channels each. scATAC-seq processing and library preparation was continued following the manufacturer’s protocol. Sequencing of the single-cell libraries was performed in the Luca/Pique-Regi laboratory using the Illumina NextSeq 500 and 75 cycles High Output Kit.

### Processing of single cell ATAC raw data

We employed `cellrange-atac (v1.2.0) count` to map the raw sequencing reads to the GRCh37 human reference genome (see data quality statistics in Table S3). After alignment and peak calling in each library separately, we used the peaks determined by cellranger to define a common set of peaks across all libraries using the `reduce` function in `GenomicRanges R` package and quantified the reads mapped to the peaks using `FeatureMatrix` function in `Signac R` package.

To assign cells to individuals, we used the `popscle` pipeline (`dsc-pileup` followed by `demuxlet`) with the default parameters (Kang et al. (2018)). The VCF file used for demuxlet contained 1,563,685 SNPs which were covered by at least 100 scATAC-seq reads across all libraries. After removing the doublets and ambiguous droplets that were detected from `demuxlet`, a total of 98,818 cells remained for downstream analysis. Each remaining droplet was assigned to one of the 16 individuals with the highest likelihood based on its genotypes. We then used these results to filter the Signac object and add the individual labels to each cell. A median of 1,186 cells per individual were obtained across five conditions (Figure S1A).

### Clustering and visualization of data

We performed clustering analysis for the scATAC-seq data using `Signac R` package Stuart et al. (2021). We used the term-frequency-inverse document frequency (TF-IDF) method to normalize data by running `RunTFIDF`. We applied singular value decomposition (SVD) on the TF-IDF matrix by running `RunSVD` (`min.cutoff=“q0”`) using all the features. This dimensional reduction method (combination of TF-IDF followed by SVD) is known as latent semantic indexing (LSI). The first LSI component was strongly correlated to sequence depth rather than biological variation. Hence the LSI components from 2nd to 50th were selected to construct a Shared Nearest Neighbor (SNN) graph using `FindNeighbors` and then cell clustering was performed using `FindClusters` with a resolution of 0.15. Finally, we constructed Uniform Manifold Approximation and Projection (UMAP) with 2-50 LSI components to visualize the clustering results.

### Integration of scRNA-seq data and cell type annotation

To integrate the scATAC-seq data with scRNA-seq data, we first inferred gene activity from chromatin accessibility using `Cicero` (Pliner et al. (2018)). Following this, we performed the standard normalization procedure on the gene activity matrix using the `NormalizeData` function with the parameter of `scale.factor` set to be the median of the total gene activity. For cell type annotation, the scATAC-seq data was integrated with the scRNA-seq data measured from the same 16 individuals previously collected(Resztak et al. (2023)). Note that for other downstream analyses that focus on gene expression changes in mean and variability we use the full set of 96 individuals with scRNA-seq. We first mapped our scRNA-seq data to the well annotated reference PMBC dataset that was wrapped in the Seurat package (Hao et al. (2021)) and annotated the cells into the following cell types including B cells, Monocytes, NK cells, T cells (CD4 T, CD8 T or other T) and DC. Next we identified anchor cells between the scATAC-seq and scRNA-seq datasets using canonical correlation analysis by running `FindTransferAnchors` with the parameter of reduction setting to `“cca”`. Cell type labels in scRNA-seq datasets were transferred to scATAC-seq datasets using the `TransferData` function with `weight.reduction=query[[“lsi”]]` and `dims=2:50` (Figure S2A and B). We used a heatmap to evaluate how well each cluster matched cell types annotation (Figure S2C). Finally we assigned the clusters (0, 1, 3, 6-10 and 12-14) to be T cells, cluster 2 to be Monocytes, cluster 4 as B cells, 5 as NK cells and cluster 11 to be DC.

In order to obtain more accurate peak definition across all libraries, we performed peak calling for each cell type using `MACS2` algorithm (Zhang et al. (2008)) by running `CallPeaks` function in `Signac` with the parameters of `group.by` set to be cell type. We then merged the peaks with the `Reduce` function as above resulting in 268,849 features and we quantified the reads mapped to peaks in each cell with a median of 6,384 features per cell across the five conditions (Figure S1, Table S4).

We annotated peaks using the `annotatePeak` function in the R package `ChIPseeker` with the distance parameter `flankDistance` set to 100 kb (Yu et al. (2015)). We observed more than half of the peaks (66.92%) fall into enhancer regions, including distal intergenic and intronic regions) and only around 28.03% of the peaks are in promoter regions (Figure S3). Through `ChIPseeker`, we also annotated the peaks to the closest genes within 100kb.

### Differential accessibility analysis

We generated pseudo-bulk ATAC-seq data by summing the reads for each feature and each sample across all cells from each of the five cell types (B cell, dendritic cell, Monocyte, NK cell and T cell), separately. A total of 260,822 features from autosomes and 331 combinations of cell-type+treatment+individual were considered to create a pseudo bulk count matrix (at least 20 cells were required for each combination and each feature has at least 20 reads mapped across all the cells). For each cell type separately, we carried out differential chromatin accessibility analysis using `R DESeq2` package (Love et al. (2014)) and the following model (`∼ individual+treatment`). We then used the following four contrasts for the treatment variable: (1) LPS, LPS vs CTRL; (2) LPS+DEX, LPS+DEX vs LPS; (3) PHA, PHA vs CTRL; (4) PHA+DEX, PHA+DEX vs PHA. The `DESeq2` fits a generalized linear model (GLM) in the statistical inference for differential analysis by assuming read counts for *i*_*th*_ gene in *j*_*th*_ sample (*K*_*ij*_) following a negative binomial distribution with mean *µ*_*ij*_ and dispersion *α*_*j*_,

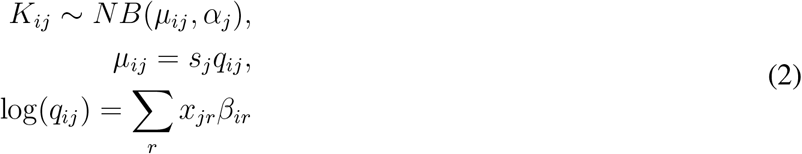

The size factor *s*_*j*_ is calculated by the median-of-ratios to account for differences in sequencing depth between samples (default settings). The *x*_*jr*_ and *β*_*jr*_ denote the treatment design matrix element and its coefficient. A prior to shrink the estimation of LFC towards zero for reducing the noise from low expressed reads is also used (default mode). Differentially accessible regions (DARs) were defined as those with FDR less than 10% and absolute estimated fold change larger than 1.41. To test if treatment preferentially increased or decreased chromatin accessibility, we performed a one sided proportion test using the `prop.test` R function.

To provide a biological insight into chromatin response changes, we annotated each DAR to the closest gene within 100 kb and then performed `GO` enrichment analysis using the Fisher’s exact test as implemented in the `compareCluster` function in R package `clusterProfiler` v4.0 (Wu et al. (2021)). DARs for each contrast and sign (opening and closing chromatin accessibility) were analyzed separately.

Previously we identified a total of 6,571 differentially expressed genes (DEGs) in the 96 individuals with the same contrasts and cell types as used in this study(Resztak et al. (2023)). We conducted Fisher’s exact test using the R function `fisher.test` to examine if these DEGs are enriched in DARs under the same contrast. We counted how many genes with DARs are identified as DEGs. We also applied the same approach to examine if differentially variable genes (DVGs) observed in our previous scRNA-seq study (Resztak et al. (2023)) are enriched nearby DARs. The combinations (cell type+contrast) with at least 100 DARs were considered in this enrichment analysis.

To calculate Pearson’s correlation between gene expression and chromatin accessibility responses, we first defined gene-level chromatin accessibility changes. To this end, we considered all peaks that were assigned to a gene as described above, and calculated the median LFC. The LFC values of transcriptional levels and variability were obtained from our previous scRNA-seq study (Resztak et al. (2023)).

### Motif analysis

We identified the motifs instances across all peaks using the `AddMotifs` function in the R package `Signac` with the reference genome setting to `BSgenome.Hsapiens.1000genomes.hs37d5` and the position weight matrices (PWMs) of 633 core motifs downloaded from JASPAR2020. We performed motif enrichment analysis to find which motifs are overrepresented in the DARs that were detected for each treatment contrast and direction separately by running the `FindMotifs` function in `Signac`. The background peaks of `FindMotifs` were defined as those peaks that were expressed in at least 2% of cells for each cell type. We used the same background peaks across contrast/direction in the same cell type.

We derived a motif activity value by quantifying the reads mapping to peaks with binding sites matching TF motifs using `chromVar` wrapped in the `RunChromVar` function (Schep et al. (2017)). Our motif activity value is exactly the deviation *z*-score derived in `chromVar` and is calculated in the same way as it was proposed in the original model. First the motifs in each peak are scanned with `AddMotifs` (i.e., `matchMotifs`, also used by the `chromVar` package and `motifmatchr` wrapping the `MOODs C++` library Korhonen et al. (2009)) with a default P-value cutoff of 5 × 10^−5^. Afterwards two matrices are created: i) a matrix of fragments counts in peaks *X* (rows are cells, and columns peaks), and ii) a matrix of motifs matches *M* (where rows are motifs, and columns peaks and value 1 if motif is present or 0 otherwise). Two procedures are conducted in `chromVAR` program: i) raw accessibility deviation is computed by the total accessibility of peaks with that motif minus the expected count for that cell and then divided by the expected values; and ii) the background peaks are used to correct technical biases caused by PCR amplification or variable Tn5 tagmentation (Schep et al. (2017)) using multiple background sampling iterations. Finally, a deviation *z*-score is computed by dividing the bias-corrected deviation by the s.d. of the background raw deviations.

To investigate fold changes for motif activities between treatments, we averaged the motif activity across cells for each cell-type, treatment and individual combination and then carried out differential motif activities analysis using `lm` with a linear regression model `∼ treatment` for the same four contrasts: LPS, LPS+DEX, PHA and PHA+DEX. To correct for multiple hypothesis testing, we used the `p.adjust` function with `BH` method to control for the false discovery rate (FDR) for each cell type and contrast separately. We selected the 51 TF motifs with FDR*<*10% and the response change *>*1.41 to visualize response changes across conditions in the heatmap using `Heatmap` in `ComplexHeatmap` R package (Gu et al. (2016)). The selected motifs were clustered into 4 categories using a hierarchical clustering method with `complete` linkage and `Euclidean` distance, implemented in `Heatmap`.

To model the relations between changes in TF motif activity and in TF-regulated gene expression, we defined TF-regulated genes as follows, (1) we first considered DARs for any of the four contrasts and within each cell type, and that also contained motifs; For LPS and PHA, in each cell type we considered the sign of the maximum motif activity change between the two conditions, and this indicated the direction for the response to both immune stimulants. For example, if a motif has the highest change in activity in LPS and the sign is positive, we only selected DARs containing this motif and with a positive LFC either in LPS or PHA. For LPS+DEX and PHA+DEX, we used a similar strategy to select the DARs with the same direction of change to the one of the motif activity. We then considered the union of DARs across both DEX and no-DEX conditions for the next step. (2) we considered the closest genes for the DARs with motifs identified in step 1; (3) we only used the genes that were differentially expressed (gene expression or variability) in any contrast for each cell type. We calculated the median of LFC on gene expression of these selected DEGs as a measure of target gene response. The same procedure was used to measure the gene expression variability response for target genes. Finally, we calculated Pearson’s correlation between the TF motif activity response and the TF-regulated gene expression or variability response.

### Defining a functional annotation of genetic variants

We annotated 6,472,200 genetic variants tested in the ALOFT study (MAF *>*10%, Resztak et al. (2021)) based on two components of our scATAC-seq data analysis: (1) whether these genetic variants are falling within cis regulatory elements (CRE) (ATAC peaks) for each cell type and (2) whether these genetic variants are in TFs motif instances within those peaks. For the first annotation, we defined cell type active peaks as those that were expressed in at least 2% cells for each cell-type resulting in a total of 54,783, 79,263, 52,024 and 45,462 peaks for B cell, Monocyte, NK cell and T cell, respectively. For the second annotation, we downloaded the position frequency matrices (PFM) of 633 human core TFs motifs from JASPAR2020 database (Fornes et al. (2020)) and scanned the GRCh37 human reference genome using `scanPwmVar` (Pique-Regi et al. (2011)) to determine if genetic variants are in TF motifs instances with a match score (log_2_ likelihood ratio) higher than 10 either for the reference or alternative allele. We classified these motifs into cell type active motifs and response motifs. First we obtained a set of cell type active motifs by fitting logistic regression using the least absolute shrinkage and selection operator (LASSO) model (`glmnet`) (Friedman et al. (2010)) based on the following model:

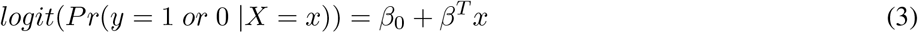

Where *y* represents the status of cell type active peak, 1 denoting activated status and 0 being inactive status; *x* being the occupancy matrix of 633 motifs for the specific ATAC peak, 1 denoting the motif occupied in specific region and 0 otherwise; and *β* being the coefficients of the TF motif contributing to cell type status. We estimated standard error of coefficients using 20 bootstraps. We defined cell type active motifs by *p <*0.05 and *β* >0. We identified 155, 168, 152 and 181 active motifs for B cell, Monocyte, NK cell and T cells, respectively. The response motifs were those identified in the differential motif activity analysis for each condition (cell type+contrast) with FDR*<*10% and the absolute changes of motif activity higher than the top 10% values. For the lymphoid cell types, we defined the top 10% changes as the threshold values across contrasts for each cell type. The threshold values are 0.26, 0.28, and 0.21 for B cells, NK cells and T cells, respectively. For the monocytes, we defined the top 10% changes as the threshold values across cell types and contrasts, the value being 0.32.

We then prepared three kinds of annotation files: (A) Cell type-specific simplified annotation files with three categories of genetic variants in a single column for each cell type, 1-**Peak** representing those genetic variants falling within cell type active peaks not hit by the cell type active motifs or response motifs (union of the four response motifs in the specific cell type), 2-**Active motif** being those genetic variants within peaks only hit by cell type active motifs not by response motifs and 3-**Response motif** being those genetic variants within peaks also hit by response motifs. The genetic variants outside of cell-type active peaks are regarded as background annotation while also controlling for distance to the TSS. This file is used for cell type-specific eQTL mapping from scRNA-seq. (B) Simplified annotation file with a single column that is used for eQTL mapping in whole blood bulk data (Resztak et al. (2021) and Aguet et al. (2020)). We still categorized the genetic variants into the above three groups while not considering cell type specificity, including **Peak, Active motif and Response motif**. (C) the condition-specific annotation file with 16 columns corresponding to 16 conditions of response motifs (4 cell type × 4 contrasts) respectively. For each column, the genetic variants within the active peaks hit by condition-specific response motifs are encoded to be 1 while other variants are treated as background annotation. We used this annotation for the two bulk eQTL mapping analyzes to estimate the enrichment levels for each condition response motifs.

### Liftover the genetic variants from human reference genome GRCh37 to GRCh38

As we had to utilize both GRCh38 and GRCh37 for some of the analyses, we had to map the studied genetic variants from GRCh37 to the GRCh38 version. We employed the `LiftOver` tool to bring the SNPs to the GRCh38 version. For the SNPs that did not have dbSNP rs number, we first looked up that information based on genomic coordinates by running the `annotate variation.pl` with `-filter` option in `ANNOVAR` package (Wang et al. (2010)). Among 6,472,200 SNPs, a total of 6,427,404 SNPs were mapped to the GRCh38 reference genome.

### Enrichment analysis of genomic annotation for eQTLs

We used a Bayesian hierarchical model implemented in `TORUS` (Wen (2016)) to estimate the enrichment level for each genomic annotation. We provided the summary statistics from eQTL studies or asthma GWAS for the `TORUS` tool. We used data from three eQTL studies, including whole blood bulk data from the ALOFT project with 251 asthmatic children (Resztak et al. (2021)), whole blood bulk data from GTEx cohort with 670 healthy individuals (Aguet et al. (2020)), and the PBMC single cell eQTL dataset (4 cell type × 5 conditions) from 96 children with asthma, a subset of ALOFT (Resztak et al. (2023)). The summary statistics of the three eQTL studies were obtained by running `FastQTL` software and testing genetic variants within 1 MB of the transcription start site (TSS) for each gene. The latter two annotation (B and C) files are used for two bulk eQTL studies while the annotation files (A) for the single cell eQTL dataset. Note that for the two simplified annotation files with single column we get the estimation of enrichment level which are relative to those SNPs that are not covered in the peaks regions, while for the condition-specific annotation file (C) we get the estimation of enrichment levels for each condition response motif that are approximately relative to the SNPs within peaks.

### Fine-mapping of eGenes using DAP-G

We performed the fine-mapping analysis for gene expression in the PBMC single cell dataset and the whole blood bulk datasets from ALOFT and GTEx V8 project using the `DAP-G`, a bayesian multi-SNP association model with a deterministic approximation of posterious (DAP) algorithm (Wen et al. (2016)). To integrate the simplified annotation derived from scATAC-seq data, we provided the SNP prior probability file for `DAP-G`, which was calculated by `TORUS`. For fine-mapping analysis, we utilized the cell type-specific simplified annotation files.

### Identification of asthma associated genes

We first conducted a TWAS analysis to identify genetically regulated gene expression correlated with asthma risk by integration of the eQTL mapping results with Asthma GWAS. We used the GWAS meta-analysis summary data for asthma disease from 18 biobanks in Global Biobank Meta-analysis Initiative(GBMI, Tsuo et al. (2022)) including 38,208,452 genetic variants in autosomes. The above two bulk eQTL mapping datasets were used in the TWAS analysis. Similar to the SMR *f*approach (Zhu et al. (2016)), we selected the top eQTL for each gene but we used two alternative criteria, 1) the minimum p-value from the `FastQTL` output, or 2) the highest PIP from the fine-mapping results obtained using `DAP-G` with response motif annotation. In some cases the top eQTL was not tested in the GWAS. In that case, we selected the closest SNP (within 10 Kb) in the GWAS. We then calculated the SMR *p*-value as previously reported (Zhu et al. (2016)) and used the `p.adjust` function with the Benjamini-Hochberg’s `BH` approach to control for the false discovery rate. The asthma associated genes by TWAS are defined as those with the FDR smaller than 10%.

We compared the results using the two approaches to select the regulatory variant for SMR. We used the fine-mapped eQTL results obtained by integrating the functional annotation and compared the results with selecting the SNP with minimum *p*-value for each eQTL signal. We obtained a greater number of genes significantly associated with asthma (Figure S24) using the fine-mapping approach both when considering the ALOFT and the GTEx whole blood data (Figure S24, Table S12 and S14).

TWAS approach has an inflation in the detection of genes linked to the specific traits due to the presence of linkage disequilibrium between eQTLs and causal GWAS variants (Hukku et al. (2022)). To remedy this issue, we also performed colocalization analysis of asthma GWAS and whole blood eQTL data using `fastENLOC` (Wen et al. (2017)), which aimed to assess the overlap of causal GWAS hits and eQTL signals. We prepared the two following files as the input of `fastENLOC`: (1) fine-mapped eQTLs from the ALOFT cohort with integration of Response motif annotation using `DAP-G` and (2) fine-mapped asthma GWAS data using `TORUS`. Via integrating the prior probability for gene expression and asthma risk, we obtained the gene locus-level colocalization probability (GLCP) for asthma risk in the colocalization analysis. TWAS and colocalization analysis have limitations and inconsistencies but their joint usage can yield robust and powerful inference results (Okamoto et al. (2023)). Using the `INTACT` R package (Okamoto et al. (2023)) we combined the colocalization results (GLCP) and TWAS *z*-score to derive the probability of putative causal genes for asthma risk. The putative causal genes are defined as those with FDR*<*10% from `INTACT`. We characterized the causal variants for these asthma risk genes using the above annotation. We also predicted the binding score of the causal variants that were bound by TFs using a position weight matrix (PWM) model and `scanPwmVar`. Following the criteria in the previous study (Moyerbrailean et al. (2016a)), we determined the genetic variants that were predicted to alter the affinity of binding TF (≥ 20 fold).

## Results

### scATAC-seq data identifies expected major clusters of immune cell types

We performed scATAC-seq using the 10x chromium platform. After quality control and filtering, we obtained high quality open chromatin profiles for 98,818 cells from 16 individuals across 5 conditions, corresponding to a median of 1,186 cells per individual and condition (Figure S1, Table S4). In the integrated scATAC-seq and scRNA-seq dataset, we identified four major cell types (B cells, Monocytes, Natural Killer (NK) cells and T cells) and also one minor cell type, Dendritic cells (DCs) (Figure 1B). In this integrated dataset, NK cells can be more clearly distinguished from T cells compared to scRNA-seq data alone (Resztak et al. (2023)). As expected, T cells formed the largest cluster with 70,824 cells (about 71%), followed by Monocytes (10,287 cells, about 10%), B cells (8,586 cells, about 8.7%), NK cells (8,037 cells, about 8.1%) and Dendritic cells (DC, 1,084 cells, 1%). We also confirmed the cell type annotation using the chromatin accessibility of canonical cell type marker genes (Figure 1C). While the clustering results were primarily dominated by cell type, we also observed cells were partially separated by treatments, especially in NK cells, which were separated into DEX treated and non-DEX treated (Figure S4). In order to identify chromatin accessibility regions, we performed peak calling for each cell type using MACS2 (Zhang et al. (2008)) and obtained a set of 268,849 candidate cis-Regulatory Elements (cCREs) across 98,818 cells.

### Changes in chromatin accessibility are associated with changes in gene expression mean and variability

To characterize chromatin accessibility changes induced by immune stimuli and glucocorticoids, we first aggregated the feature reads across cells for each combination (cell type-individual-condition) to generate pseudo-bulk data. We used DESeq2 (Love et al. (2014)) to identify differentially accessible regions (DARs) within each cell type for the following contrasts (Figure S5, S6 and Table S8), (1) LPS versus CTRL; (2) LPS+DEX versus LPS; (3) PHA versus CTRL; (4) PHA+DEX versus PHA. Across all five cell types and contrasts, we detected a total of 62,424 DARs (FDR*<*10% and |log_2_ *FC*| *>*0.5, Figure 2A). Both LPS and PHA induced the strongest chromatin response in monocytes (5.8% and 10.1% tested peaks are DARs, respectively) compared to other cell types, followed by T cells for PHA only (2.5% tested peaks are DARs) (Table S5). This reflects the underlying biology: monocytes are the immune cells that respond to bacterial infection (LPS is a component of the bacterial membrane), while T cells as expected respond strongly to PHA which is a T cell mitogen. In contrast, glucocorticoids induced a strong response in all lymphocytes (Figure 2A), especially in T cells (4.2% tested peaks are DARs for LPS+DEX and 6.7% tested peaks are DARs for PHA+DEX, Table S5).

**Figure 2.**
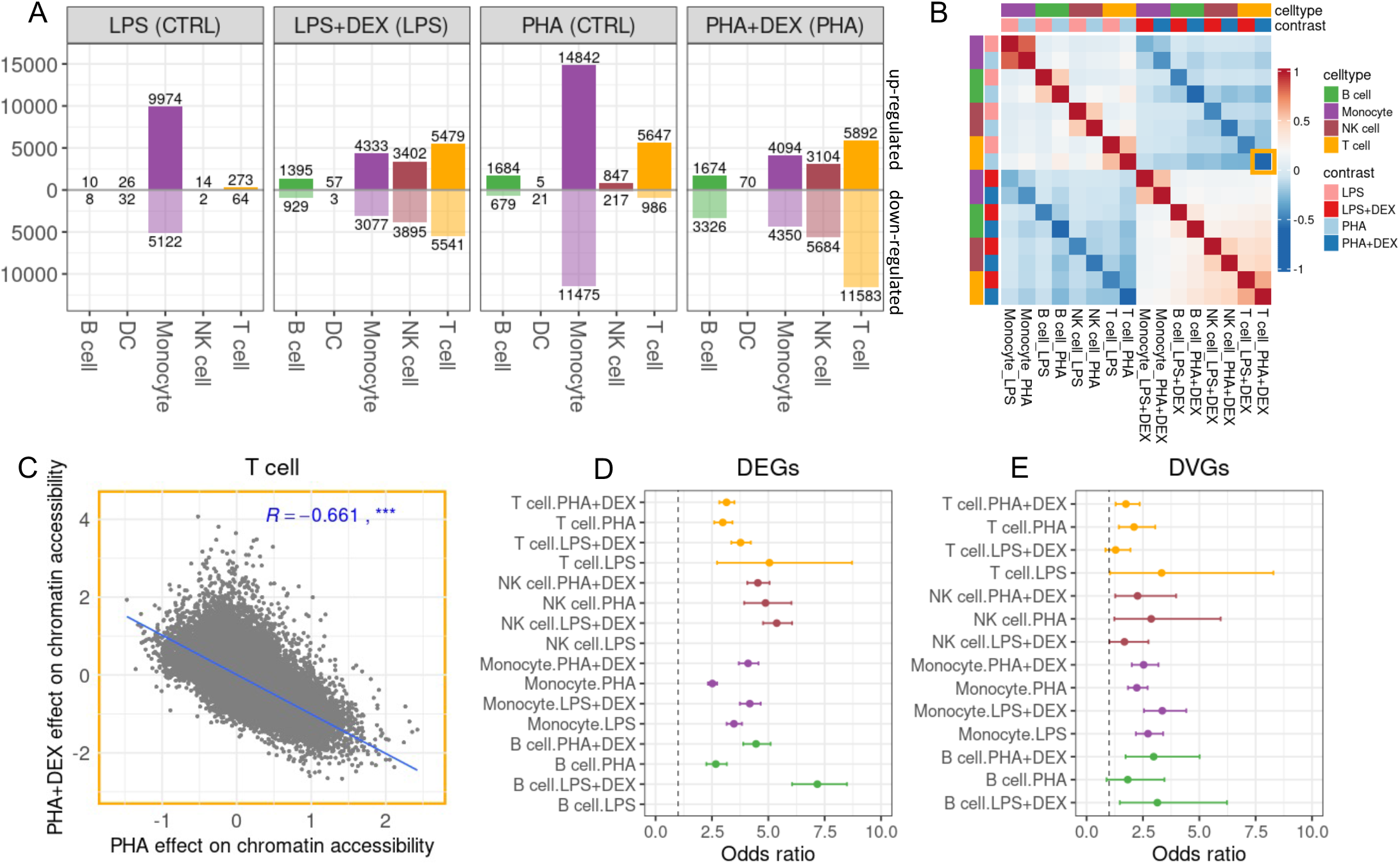
Identification of differential accessible regions (DARs) in response to immune treatments across cell types. **A** Number of DARs across cell types, B cells (green), DCs(grey), Monocytes (purple), NK cells (maroon) and T cells (orange) for LPS (LPS vs control), LPS+DEX (LPS+DEX vs LPS), PHA (PHA vs control) and PHA+DEX (PHA+DEX vs PHA). DARs are defined as those with FDR*<*0.1 and |log_2_ *FC*| *>*0.5. Below axis represents closing chromatin regions while above axis being open chromatin regions. **B** Heatmap of pairwise Pearson correlation of log_2_ fold change(LFC) of union of 62,424 DARs across 16 conditions, 4 major cell types (excluding DC cells) and 4 contrasts, red color corresponding to stronger positive correlation values while blue color for negative correlation values. **C** The scatterplot of the effects of PHA+DEX on the 62,424 DARs (y-axis) against those of PHA on chromatin accessibility (x-axis) in T cells, Pearson correlation coefficient R=-0.661 (***, *p*< 0.001). Blue line represents the regression relationship of PHA+DEX effects against PHA effects. **C and D** Forest plots of the odds ratios that differentially expressed genes (DEGs) or differentially variable genes (DVGs) are enriched in DARs, respectively.

When considering whether the treatments result in increased or decreased chromatin accessibility, we observe 2-fold (*p <* 2.2 × 10^*-*16^) increase in accessibility induced by LPS in monocytes (Table S6). In T cells we clearly observe the antagonistic effects of immune-stimulation and glucocorticoids, with 6-fold (*p <* 2.2 × 10^−16^) increase in accessibility induced by PHA and 2-fold (*p*< 2.2 × 10^−16^) decrease in accessibility induced by DEX (Table S6).

Overall, we observed that effect of dexamethasone and the immune stimuli on chromatin accessibility were negatively correlated (Figure 2B and C, Figure S7), which implicated that glucocorticoids prevents the chromatin accessibility changes induced by immune stimuli (the same pattern as observed in transcriptional changes by Resztak et al. (2023)). This result confirmed the role of chromatin accessibility changes in mediating the immune response (Pacis et al. (2015)). Indeed, genes nearby DARs were significantly enriched (FDR*<*10%) for immune-related pathways, such as type I interferon signaling and response to LPS (Figure S8).

To investigate to what degree changes in chromatin accessibility result in transcriptional changes in response to immune treatments, we considered both changes in gene expression mean and variability. We found that the number of DARs was correlated with the number of differentially expressed (DEG) and differentially variable (DVG) genes (Figure S9). However, when we consider individual genes, we find significant and positive correlations only between changes in chromatin accessibility (DAR LFC) and changes in gene expression mean (DEG LFC, Figure S10), while correlations are not significant with changes in gene expression variability (DVG LFC, Figure S11). To further investigate changes in chromatin accessibility as a main mechanism leading to transcriptional response, we considered DARs near DEGs and DVGs. We identified 81.7% DEGs and 75.6% DVGs within 100 Kb of a DAR, corresponding to a significant enrichment for all cell type-treatment combinations for DEGs (2.5–7.22 fold enrichment, *p <* 1 × 10^−5^, Figure 2D). For DVGs, the enrichments were more moderate and remained strongly significant in Monocytes (Figure 2E) with a fold enrichment of 2.4–3.7.

### Increase of TF binding increases the mean and tightens the variability of gene expression

The concordance observed between chromatin and gene expression changes can be explained by the hypothesis that environmental factors, such as immune treatments, induce the recruitment of specific transcription factors (TFs) to DNA sequences in accessible regions, thus triggering transcriptional changes of target genes. To test this hypothesis, we derived a motif activity value that quantifies the reads mapping to peaks with binding sites matching TF motifs using `chromVAR` (Schep et al. (2017)). Intuitively, these motif activities can be interpreted as a genome-wide measurement of transcription factor binding activity to their target sequence. Reassuringly, the motif with one of the highest increased activity after treatment by glucocorticoids is the NR3C1 motif (Figure 3A), which also displayed a high enrichment in DARs in the DEX conditions across cell types (*p <* 2.2 × 10^−16^, Figure S12). *NR3C1* encodes the glucocorticoid receptor (GR), which is activated upon glucocorticoid binding (Escoter-Torres et al. (2019)). When considering all the treatments and the motifs with largest changes in activity we can see that they cluster in four major patterns (Figure 3A): the glucocorticoid receptor cluster (GR cluster), the CCAAT enhancer-binding proteins cluster (CEBP cluster), the AP-1/NFKB cluster, which is the largest and most heterogeneous, and the STAT/IRF cluster. These factors are all known to interact with the GR to regulate gene expression (Boruk et al. (1998); Nissen Yamamoto (2000); Biddie et al. (2011)). The CEBP cluster is specific to monocytes and is activated by both immune-stimuli (LPS, PHA), while its activity remains unchanged by glucocorticoid treatment. The AP-1/NFKB cluster is also largely specific to monocytes, except for NFKB/RELA motifs. This cluster is also activated by the immune-stimuli, but glucocorticoids reduce its activity in a cell type and TF-specific fashion. This is in contrast to the STAT/IRF cluster that is activated by the immune-stimuli (mostly PHA) and inactivated by glucocorticoids across all cell types.

**Figure 3.**
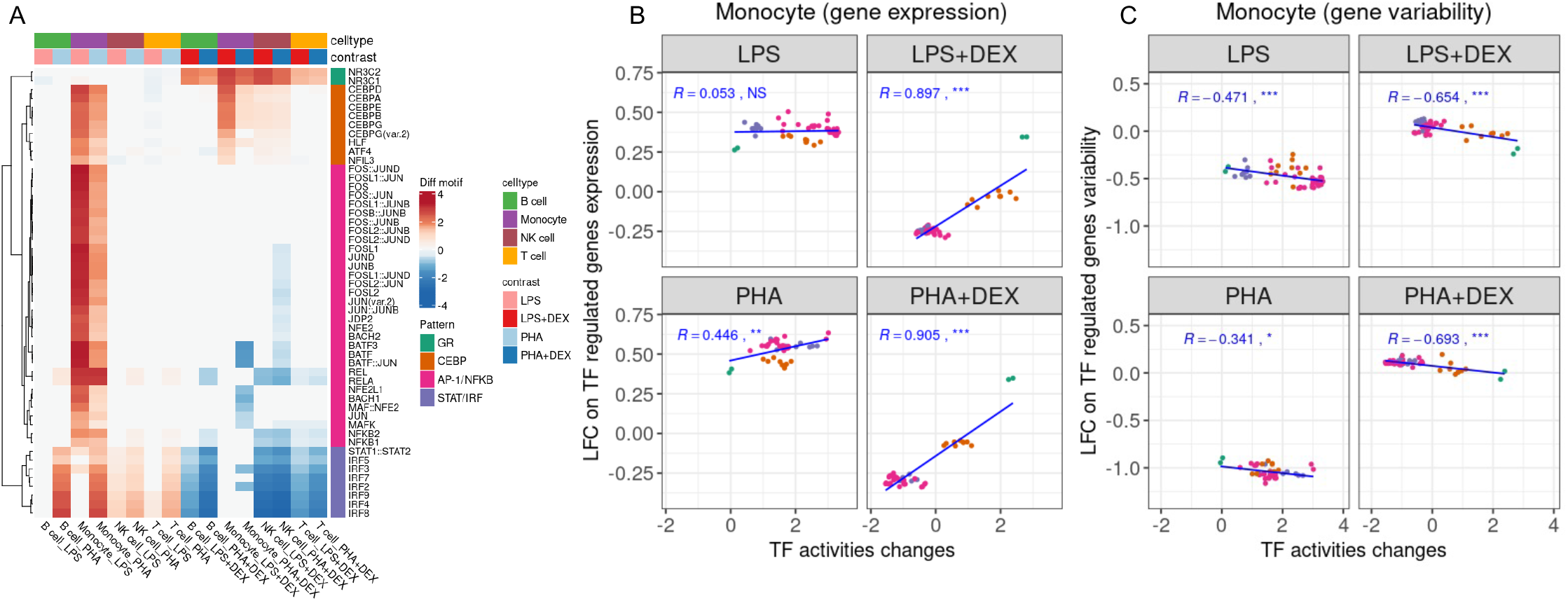
Characterization of the transcription factors motif activities induced by immune treatment. **A** Heatmap of the relative changes of motif activities in response to immune treatment for 51 TF. The color reflects the magnitude of motif activity change (red for increased changes and blue for decreased changes). Four distinct patterns of motifs are identified and colored by different colors. **B-C** The scatter plots of changes of TF motif regulated gene expression (**B**) and TF motif regulated gene variability (**C**) represented by Y-axis against those of TF motif activities (X-axis) in the monocyte in the four contrasts, LPS, LPS+DEX, PHA and PHA+DEX. R value represents Pearson correlation coefficient, NS not significant, * *p*< 0.05, ** *p*< 0.01 and *** *p*< 0.001. Blue lines represent the regression between changes of TF motif regulated gene expression (variability) against changes of TF motif activities.

We then explored the extent to which transcriptional responses were influenced by changes in TF activities. For each TF, we considered the motif activity and the downstream changes in the expression of target genes, and observed a highly positive correlation across cell types (Figure 3B, S13, S14, S15 and S16). The largest correlation is observed in the dexamethasone-treated Monocytes (Pearson correlation = 0.9 (Figure 3B)); it is largely driven by the GR, the IRF, and the AP1-NFKB clusters. This result confirms the two main mechanisms for the anti-inflammatory effect of glucocorticoids through activation of the GR: 1) binding of activated GR to glucocorticoid response elements (GRE), thus inducing antiinflammatory gene expression (Buckingham (2006)), and 2) repression of inflammatory gene expression via antagonistic interaction with other TFs (e.g. RELA and NFKB1), which prevents their binding to target genes, Escoter-Torres et al. (2019)).

We also investigated the effects of transcription factor binding on their target genes’ transcriptional variability. The increase of the motif activity is generally negatively correlated with gene expression variability (Figure 3C, S13, S17, S18 and S19). Note that this measure of gene expression variability has been already corrected by the mean-variance dependence. This result suggests that an increase of TF binding tightens the variability of gene expression around the mean. All different motif clusters observed here seem to contribute to this phenomenon.

### eQTLs in GTEx and an asthma cohort are enriched in context-dependent regulatory elements

The main mechanism underlying eQTLs is disruption of transcription factor binding to cis-regulatory elements (Claussnitzer et al. (2020)). The regulatory region activity can be modulated by the cellular context, thus resulting in cell type or treatment-specific genetic regulation of gene expression. To investigate this context-dependent regulatory activity, we derived an annotation for genetic variants in chromatin accessibility peaks as follows (Figure 4A): genetic variants in response motifs (Response motif), genetic variants in active motifs that are not response motifs (Active motif), genetic variants in all remaining peaks (Peak). We performed the enrichment analysis of these three genomic categories for eQTL obtained from the scRNA-seq data using `TORUS` (Wen (2016)) and subsequently applied the resulting enrichment information to aid eQTL fine-mapping analysis using `DAP-G` (Wen et al. (2016)). We focused on T cells because they are the most abundant cell subpopulation and the one with the largest number of eQTLs. Across the five different treatment conditions, eQTL signals were enriched for all three annotation categories with a consistently greater enrichment for Response motifs (Figure 4B). We fine-mapped a total of 519 eGenes (FDR*<*10%) associated with at least one genetic variant (Figure S21, Table S7, Table S9), which is a greater number of eGenes compared to using `TORUS` (385 eGenes) and `FastQTL` with 1000 permutation (343 eGenes) at the same significance level (FDR*<*10%). Note that with methods such as MashR that can share effect size information across cell types, a higher number of eGenes can be detected; but here we focus on the response annotations and ability to fine map and characterize eQTL loci and not eGenes.

**Figure 4.**
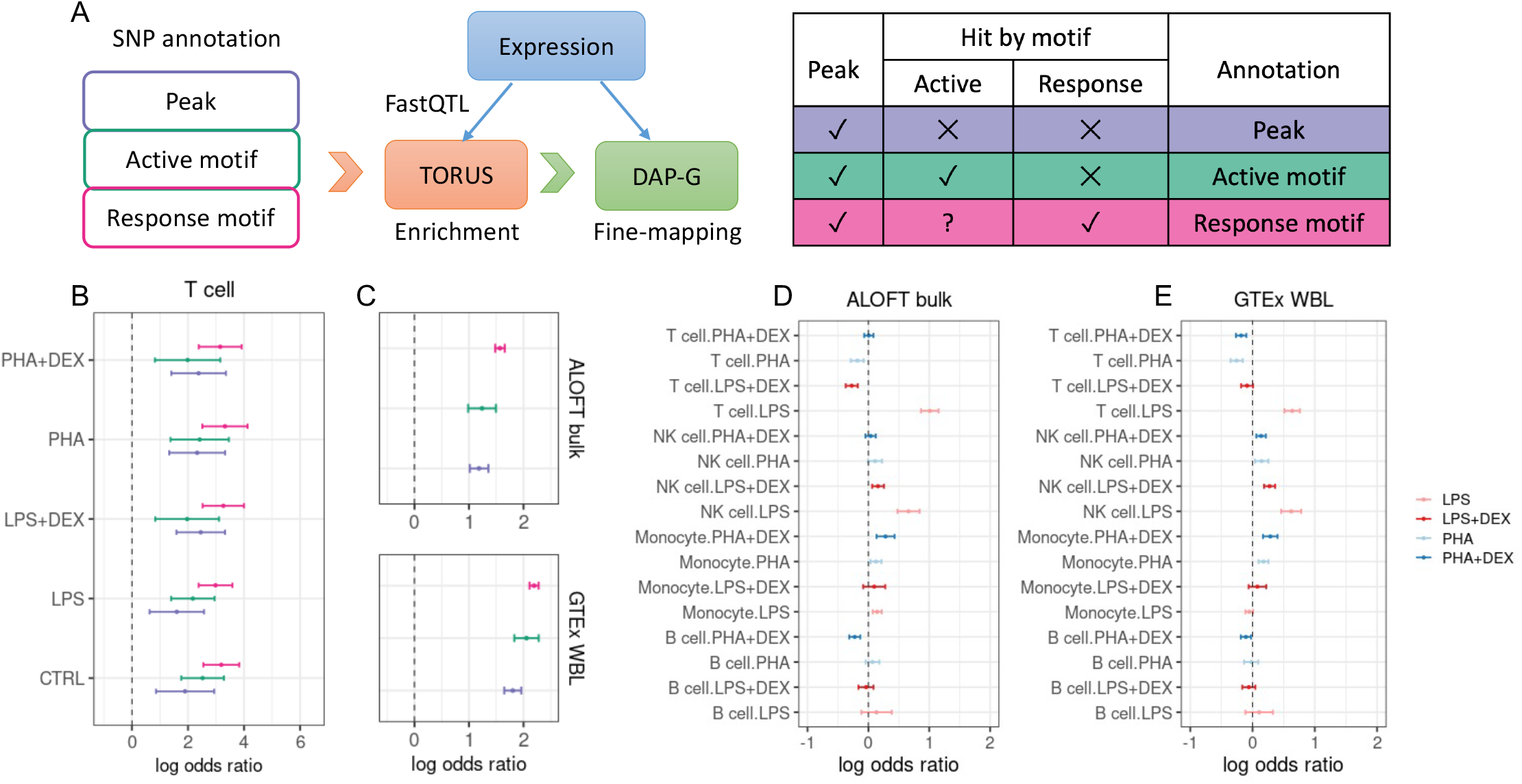
Interpretation of the regulatory mechanism underlying genetic effects on gene expression obtained from eQTL analyses. **A** Diagram of genetic analysis for gene expression and annotation of regulatory variants in peaks and if they are additionally on active motifs that change activity with the treatment (Peak, Active motif and Response motif). Using the eQTLs summary data from FastQTL, we performed enrichment analysis for three categories of variants by running TORUS. We employed DAP-G to fine-map eGenes by integration of prior information provided by TORUS. **B** Forest plot of the log odds ratio of three genomic categories enriched in five conditions single-cell-eQTL datasets in T cells. **C** Forest plot of the log odds ratio of three genomic categories enriched in two Bulk eQTL datasets of whole blood, ALOFT (top) and GTEx (bottom). **D-E** Forest plots of the log odds ratio of 16 condition Response motifs enriched in ALOFT and GTEx, respectively.

Existing population cohorts for genetic studies of gene expression include individuals that may have an arbitrary immunomodulatory state resulting from different endogenous cortisol levels or exogenous inflammatory agents that can be considered latent exposures. We have previously shown that annotation of response regulatory variants can identify eQTLs that are modulated by an environmental factor (Findley et al. (2019, 2021)). To examine the contribution of latent pro- and anti-inflammatory latent contexts to the genetic regulation of gene expression in peripheral blood, we performed enrichment analysis for the three genomic features in two bulk eQTL datasets of whole blood: the Asthma in the Lives of Families Today (ALOFT) cohort (16 samples overlapping with the current study) (Resztak et al. (2021)), and GTEx (Aguet et al. (2020), a completely independent cohort). We observed very similar enrichment patterns for the two cohorts with a significantly larger enrichment for the Response Motif category, compared to the Peak category (Figure 4C).

As the Response Motif category is composed of 16 conditions (4 cell types × 4 contrasts), we can further dissect the latent environment by testing for an enrichment for each of the 16 categories (Figure 4D and E). In both bulk datasets, we observed the highest enrichment for T cell-LPS (log odds ratio 1.009 ±0.073 for ALOFT and 0.636 ±0.063 for GTEx) and NK cell-LPS (log odds ratio 0.659 ±0.093 for ALOFT and 0.620 ±0.083) Response Motifs. We also observed significant enrichment for other categories, including NK cell-LPS+DEX Response Motifs (log odds ratio 0.158 ±0.049 for ALOFT and 0.273 ±0.045 for GTEx), Monocytes PHA+DEX (log odds ratio 0.281 ±0.076 for ALOFT and 0.285 ±0.060 for GTEx) and Monocyte-PHA (log odds 0.125 ±0.046 for ALOFT and 0.175 ±0.040) Response Motifs. For the ALOFT whole blood bulk eQTL mapping using DAP-G with integration of our annotation, we fine-mapped 6,884 protein coding eGenes (FDR*<*10%) with a median of 1 independent eQTL signal for each gene (Figure S22A and Table S10). Among them, we also observed 983 eGenes (14% percent) regulated by multiple independent eQTLs (at least 2 eQTLs).In the GTEx whole blood bulk data, we identified 9,853 eGenes (FDR*<*10%) with a median of 1.1 eQTLs for each gene (Figure S22B and Table S11), of which 2,359 eGenes (nearly 24%) were regulated by multiple independent eQTLs. The greater discovery of eGenes and eQTLs is attributed to larger sample size in the GTEx (670) compared to 251 individuals in the ALOFT cohort. The majority of eGenes (5,837) however are shared between the two cohorts (Figure S23).

### Context-dependent annotations increase the number of asthma risk genes and provide putative molecular mechanisms

To dissect the contribution to asthma risk of pro- and anti-inflammatory mechanisms that are genetically regulated, we integrated our fine-mapped eQTLs with GWAS results for asthma from Global Biobank Meta-analysis Initiative (GBMI, Tsuo et al. (2022)) using INTACT (Okamoto et al. (2023)) (Figure 5A). Transcriptome-wide association studies (TWAS) and colocalization analysis have limitations and inconsistencies but their joint usage can yield robust and powerful inference results (Okamoto et al. (2023)). We previously demonstrated that joint TWAS and locus-level colocalization analysis improves specificity and sensitivity for implicating biologically relevant genes. Here we also used our annotation to prioritize eQTLs in the TWAS step, resulting in 36% increase in the number of genes associated with asthma, compared to a standard SMR approach both when considering the ALOFT and the GTEx whole blood data (Figure 5B, S24). This difference is even more striking when integrating colocalization and TWAS evidence using `INTACT`: we refined a total of 410 putative causal genes (PCGs with FDR*<*10%) for asthma risk (Figure 5C and D, S25, Table S12, S13), compared to 298 when not using the annotation. The additional genes found with the annotation are not enriched in any specific process or pathways, compared to the overall set of 410 genes. Of the putative 410 causal genes identified, 191 (47%) were genetically controlled by regulatory variants in Response motifs (Table S14 and S15). Of these 191 putative causal variants in Response motifs, 141 are predicted to alter TF binding (absolute differential PWM score *>*4.33). Furthermore, we observed that asthma risk genes are enriched in genes differentially expressed in response to PHA and PHA+DEX in all cell types, but we don’t find a significant enrichment for genes that respond to LPS (Figure S26). This result suggests that alteration of expression for asthma risk genes in response to altered T-cell activation, is ubiquitous across all major cell types.

**Figure 5.**
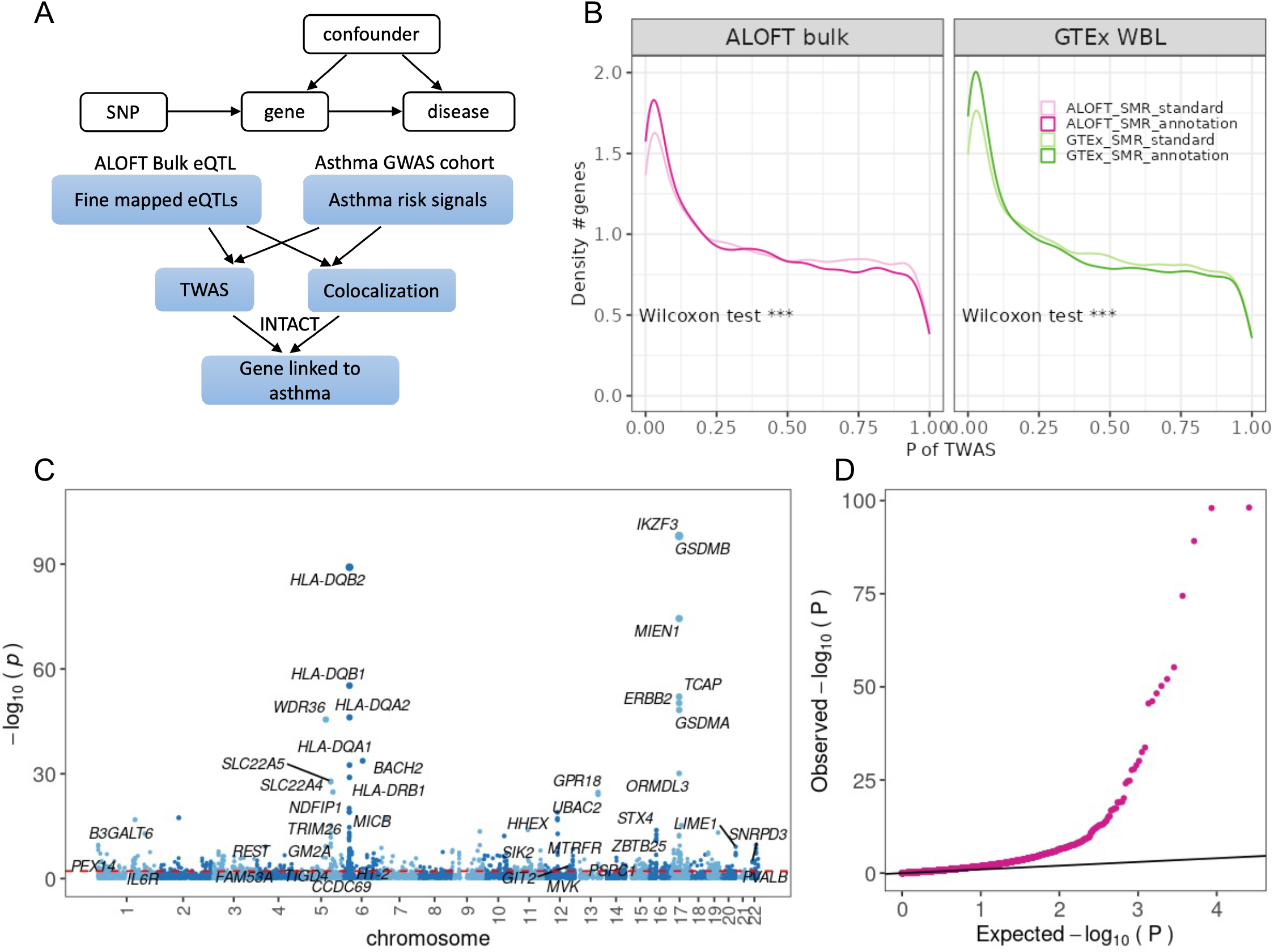
Identification of asthma risk gene by INTACT. **A** Diagram of the utilizing INTACT with combination of TWAS and colocalization to identify asthma risk genes by integration of fine-mapped eQTLs in ALOFT cohort with asthma GWAS. **B** Density plots of the TWAS *p*-value for the gene associated with asthma risk from ALOFT cohort (Left) and GTEx cohort (Right) using different approaches: light color denoting the standard SMR and dark color denoting a variant of SMR with the integration of genetic variant annotation to identify the putative causal variant for each gene **C** Manhattan plot visualization of genes genetically associated with asthma risk with TWAS with the annotation from the ALOFT cohort, each dot representing the SMR − log_10_(*p*) of each gene in the y-axis. **D** Quantile-quantile (QQ) plot for TWAS with the annotation from the ALOFT cohort.

For example, the gene *DEF6* in chromosome 6 is associated with asthma risk (FDR=7.34 × 10^−9^ from TWAS and 0.24 colocalization probability, Figure 6A). *DEF6*, known as IRF4 binding protein, encodes a guanine nucleotide exchange factor highly expressed in immune cell types which acts in regulation of immune homeostasis (Serwas et al. (2019)). Previous studies have implicated the role of this gene in autoimmune and inflammatory disorders (Binder et al. (2017); Serwas et al. (2019); Fournier et al. (2021)). We fine-mapped a SNP (rs9469982 with 0.28 PIP) genetically controlling the expression level of *DEF6*. This genetic variant falls in a peak (“6-35267195-35267857”) that is bound by a response TF (SOX8). The alternative allele of this variant has a binding score (12.47 log_2_ PWM units) for TF SOX8 that is 3 orders of magnitude higher than the reference allele (0.64 log_2_ PWM units). SOX8 shows increased activity in LPS+DEX in T cell (FDR=4.67 × 10^−12^). Meanwhile we also observed increased chromatin accessibility for this peak (6-35267195-35267857) in LPS+DEX treatment in T cell (LFC=0.283, FDR=0.026, Figure 6A).

**Figure 6.**
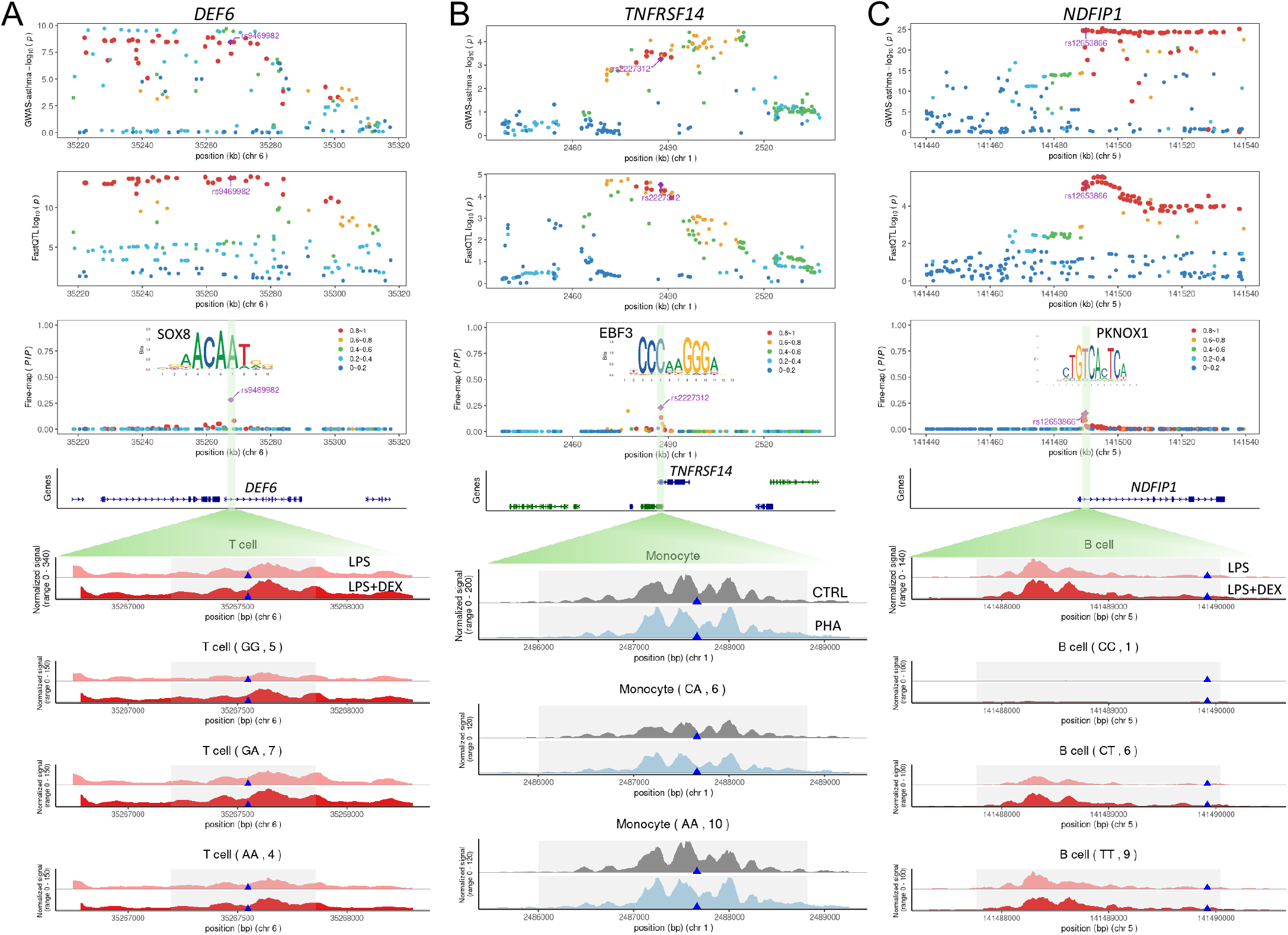
Examples of LocusZoom plots depicting candidate asthma risk genes A) *DEF6*, B) *TNFRSF14* and C) *NDFIP1*. The first four rows of plots represent: 1) *-* log_10_(*p*) of SNPs associated with asthma via GWAS and, 2) the − log_10_(*p*) for association with gene expression of the gene (eQTL), 3) posterior inclusion probability (PIP) of the SNPs being the causal eQTL using the annotation, and 4) the gene track around the lead variant. Diamond denotes the lead variant and colors representing the degree of the individual variants’ LD with the lead variant. The next following rows include normalized chromatin accessibility of CREs around the lead variant in specific cell types in contrast and treatment conditions for all the individuals. If available the individuals are split into homozygous reference allele, heterozygote and homozygous alternative allele. Note that for rs12653866/TNFRSF14 we do not have genotyped individuals for the homozygous reference.

Another example is the gene *TNFRSF14* in chromosome 1 (Figure 6B), encoding a member of the tumor necrosis factor (TNF) receptor superfamily. The protein is a key player for activating inflammatory and inhibitory immune response. This gene has also been implicated in multiple autoimmune diseases, such as in inflammatory bowel disease (Jostins et al. (2012)), ulcerative colitis (Anderson et al. (2011)) and rheumatoid arthritis (Raychaudhuri et al. (2008); Stahl et al. (2010)). The colocalization probability with asthma risk is 0.27. We identified a cluster of 25 SNPs associated with the *TNFRSF14* gene expression (signal-level PIP, 0.97), the maximum of which is the variant rs2227312 with 0.23 PIP. This genetic variant is located within a region with increased accessibility in response to PHA treatment in monocytes (1-2486002-2488823 with LFC=0.205, FDR=0.026, Figure 6B). This SNP is in a motif for the response TF (EBF3), which also shows higher genomewide activity in response to PHA treatment in monocytes (FDR=0.004). The reference allele of this SNP has a binding score for the EBF3 that is about 63 times higher than the alternative allele.

We also identified the gene *NDFIP1* as potentially associated with asthma risk (TWAS FDR=1.27 × 10^−22^ and 0.97 colocalization probability, Figure 6C). *NDFIP1* encodes a transmembrane protein that plays key roles in regulating the differentiation and maturation of T cells (Wagle et al. (2018); Altin et al. (2014); Layman et al. (2017)). Previous studies have revealed that this gene is implicated in asthma and multiple autoimmune diseases (Wan et al. (2012); Pickrell et al. (2016)). We fine-mapped a cluster of 32 variants with 0.97 joint signal-level PIP associated with the expression of *NDFIP1* (Figure 6C), the maximum of which is the genetic variant rs12653866 with 0.15 PIP. This variant is in a peak that contains a response factor (PKNOX1 or an E-box binding TF), with predicted differential allelic binding affinity. Specifically, the alternative T allele has a binding score 15 times higher than C, the reference one. In B cells, we see greater accessibility at this peak in LPS+DEX treatment (LFC=0.38 and FDR=0.02), especially in the individuals with the T allele (Figure 6C).

## Discussion

In this study, we characterized changes in genome-wide chromatin accessibility in immune cells in response to pro- and anti-inflammatory treatments at single cell resolution to study cell type specific regulation of gene expression. We show that changes in chromatin accessibility are a major driver of the transcriptional response to immuno-modulators, when considering both gene expression mean and variability. The gene regulatory response to glucocorticoids is very well characterized and known to be mediated by changes in TF binding. Here, we show that these inducible TF binding responses can be captured by studying chromatin accessibility. Furthermore, we annotated genetic variants in chromatin accessibility peaks and TF binding sites for each cell type and factors that drive the chromatin and gene expression changes. The annotation is useful not only to study and fine map single cell eQTLs, but also previous large scale bulk studies collected in unknown/uncharacterized immuno-modulatory conditions. Indeed, we show that genetic variants annotated in response factors are enriched for eQTLs both in GTEx and in a dataset of bulk RNA-seq in blood from a cohort of children with asthma, thus capturing latent inflammatory states. Overall these results highlight the potential of an approach that combines chromatin accessibility data from a relatively small sample with large eQTL and GWAS datasets. With the underlying assumption that asthma risk and asthma severity share similar biological pathways and mechanisms, at least in part, we show that the fine-mapped eQTLs are useful to further interpret GWAS signals for asthma using colocalization and TWAS techniques.

Glucocorticoids main mechanisms of action is through ligand-mediated activation of the GR, which is normally sequestered in the cytoplasm. Upon activation the GR translocates to the nucleus and binds the chromatin at response elements (GRE) resulting in regulation of target gene expression (Buckingham (2006). This gene regulatory response is achieved through cooperation with other transcription factors which modulate GR binding activity (Buckingham (2006)) and contribute to cell type specificity. In immune cells, glucocorticoids mainly exert an anti-inflammatory function, by preventing an extreme immune response which could be deadly;for example: sepsis and cytokine storms. This anti-inflammatory response is largely realized through inhibition of NF-kappaB and other pro-inflammatory transcription factors (Buckingham (2006)). Overall here we characterized chromatin accessibility at TF binding sites in asthma, thus identifying active regulatory elements in the disease status. Future studies comparing patients and controls, will be important to further dissect which of these regulatory elements and factors are specifically disrupted in the disease condition compared to healthy children.

Single cell genomics has enabled us to study gene expression levels beyond average mean gene expression and to show that even closely clustered cells have variable levels of gene expression. The physiological relevance of these transcriptional phenotypes and their role in complex traits is still under investigation. We have previously described variability in gene expression response to immuno-modulators, and demonstrated that similar to gene expression mean, gene expression variability is also under environmental and genetic control (Resztak et al. (2023)). Here we analyzed chromatin accessibility data to investigate the regulatory architecture underlying these transcriptional phenotypes. We observed that changes in motif activities are associated with changes in both gene expression mean and variability, for the same motifs. We observed that a change in motif activity had a proportional increase in gene expression mean of their target genes; while the same factors led to a proportional decrease in gene expression variability on the same target genes. Overall the same transcription factors regulate both gene expression mean and variability response. This is true both when considering a treatment that activates one main TF (dexamethasone) but also for treatments that activate a larger number of TFs (LPS and PHA). While increase in motif activity is generally associated with increase in gene expression, we observed the opposite effect with variability, where the increase of the motif activity is generally associated with a decrease in variability. It therefore appears that gene expression variability shares the same regulatory architecture as gene expression and that changes in variability result from variation in TF binding.

Environmental effects on gene regulation are pervasive even when a specific environment is not measured or accounted for. We previously demonstrated that transcriptional profiles from population-level samples bear a signature of latent environmental exposures that can be uncovered by using a context-aware annotation (Findley et al. (2019)). For example, we were able to identify a signature of caffeine exposure in the GTEx samples, by using an annotation from cells exposed to caffeine (Findley et al. (2019)). The concept of latent environment also enables the study of context eQTLs or GxE based on what is captured in the overall gene expression patterns; yet the lack of labeled environmental data or annotations reduces interpretability. Our study shows that even if we don’t have eQTL data in different contexts, we can use a context-aware annotation to reveal latent environments and increase our ability to identify risk loci that are context dependent. Importantly, we demonstrate that using a context-aware annotation increases the ability to fine-map GWAS signals for a disease with a strong GxE component. Our study thus expands recent efforts to fine-map asthma risk loci and include diverse study cohorts (Washington III et al. (2022); Tsuo et al. (2022)) by both focusing on a cohort of African American children, and considering immuno-modulatory contexts. Our study is limited in the chromatin accessibility sample, which does not allow to capture the heterogeneity of the disease in the annotation, and in the number of contexts that we were able to investigate. Despite these limitations, we provide a general framework to combine existing large genetic datasets with an experimentally generated annotation to perform GWAS fine-mapping accounting for the increasing importance of GxE in the genetic architecture of complex traits.

## Acknowledgments

This research has been supported by NIH grants from NHLBI (R01HL114097 to SZ and RS, 1R01HL162574 to FL, RP, and SZ) and NIGMS (R01GM109215 to FL and RP). We thank members of the Luca, Pique-Regi, Slatcher and Zilioli labs for helpful discussions and comments. We also thank the anonymous reviewers for comments and suggestions that helped to improve the quality of this manuscript. Finally, we also greatly appreciate the ALOFT study participants and their families.

## Author contributions

R.P-R. and F.L. designed and supervised this study. R.S. and S.Z. supervised the original ALOFT study that recruited the participants. J.R., A.A. and H.M.M. conducted the single cell experiments. J.W. performed the data analyses with the supervision of F.L., RP-R and X.W. J.W., A.R., R.P-R. and F.L. interpreted the results and led the writing of the manuscript. All the authors read, contributed edits and approved the manuscript.

## Data and Code Availability

All raw and processed sequencing data generated in this study have been submitted the NCBI database of Genotypes and Phenotypes (dbGaP; https://www.ncbi.nlm.nih.gov/gap/) under accession number phs002182.v3.p1. Large supplementary tables have been deposited in: http://genome.grid.wayne.edu/scATAC_JW/Supp_tables/. Code and scripts used to make the results are available at https://github.com/piquelab/sc-atac.

## Supplementary Figures

**Figure S1:**
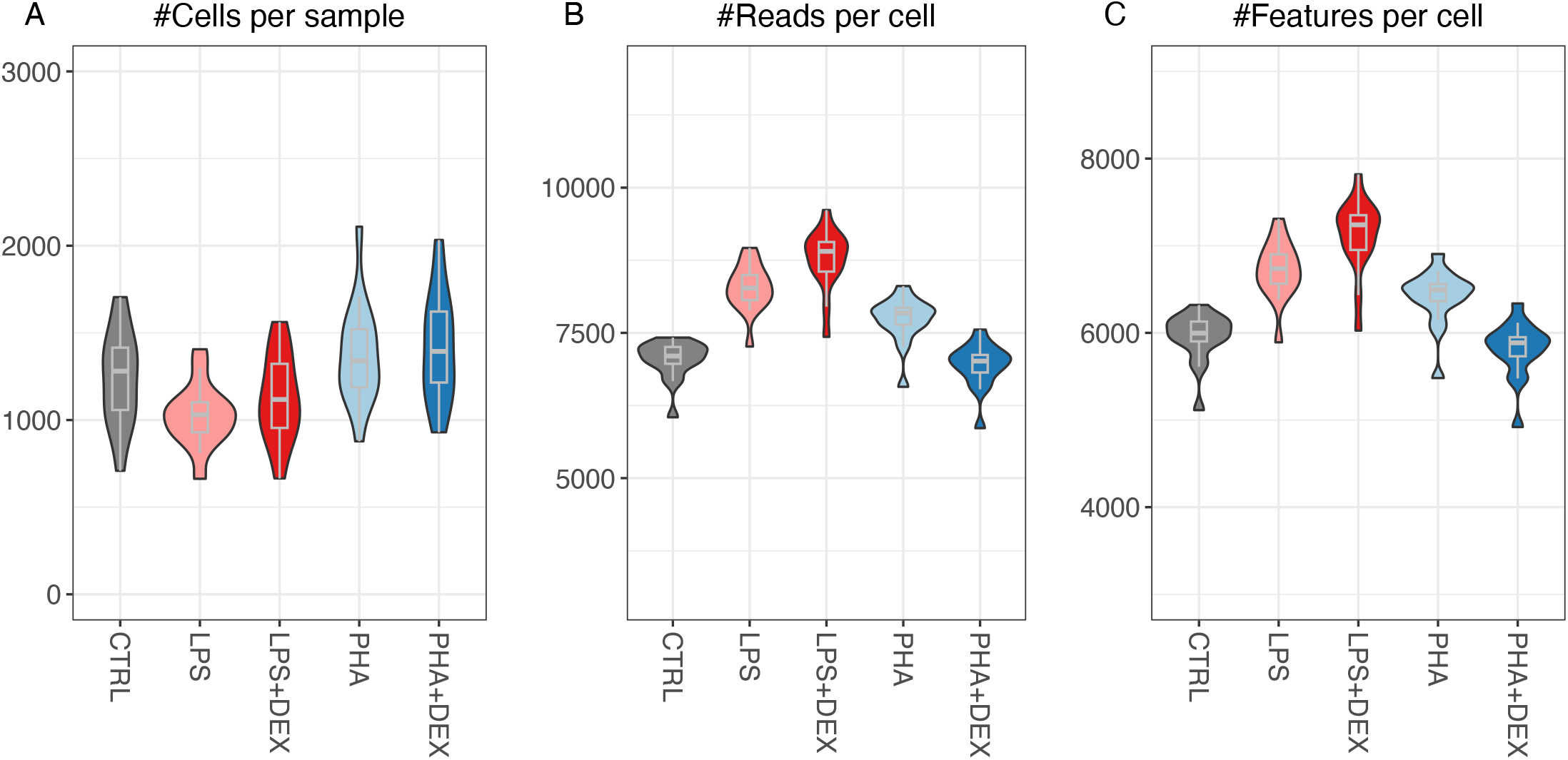
Violin plots of summary statistics per individual across five conditions. **(A)** number of measured cells per individual for each treatment. **(B)** average reads per cell from one individual for each treatment. **(C)** average number of the peaks per cell from one individual for each treatment.

**Figure S2:**
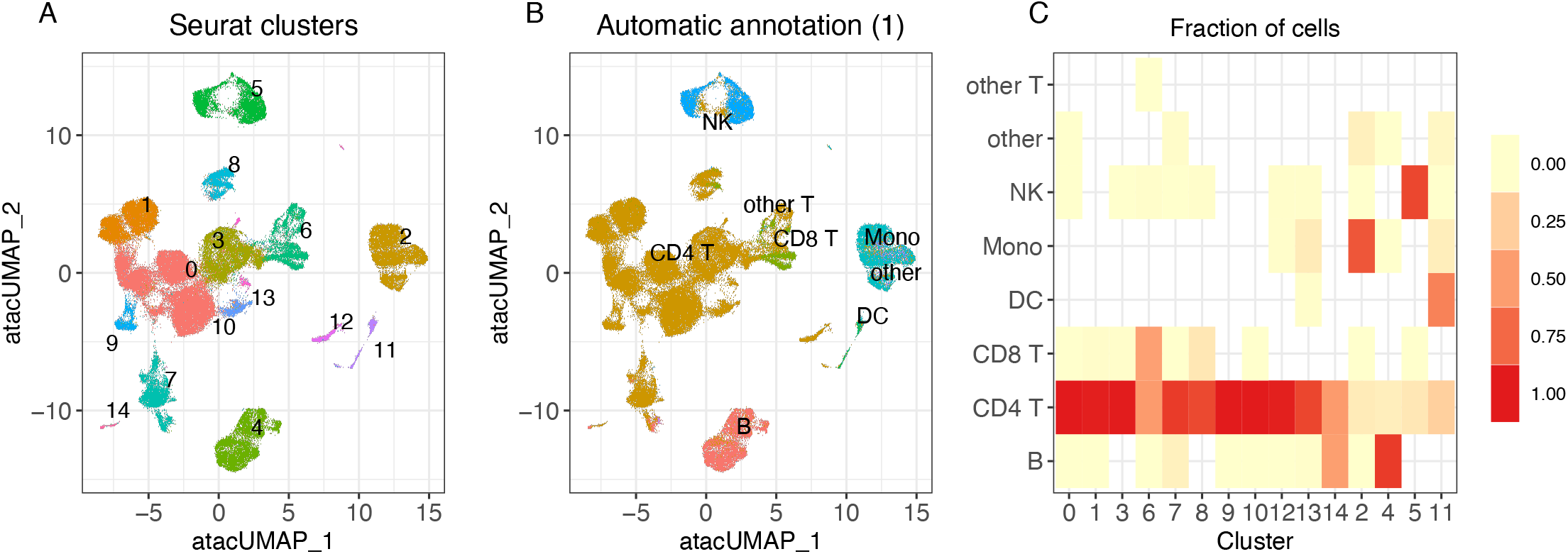
Comparison of clusters results with automatic annotation of the major cell types from the well annotated reference PBMCs. **(A)** UMAP visualization of scATAC-seq data colored by clusters **(B)** UMAP visualization of scATAC-seq data colored by the major cell types automatically annotated from the reference data. **(C)** Heatmap showing the percentage of cell types for each cluster.

**Figure S3:**
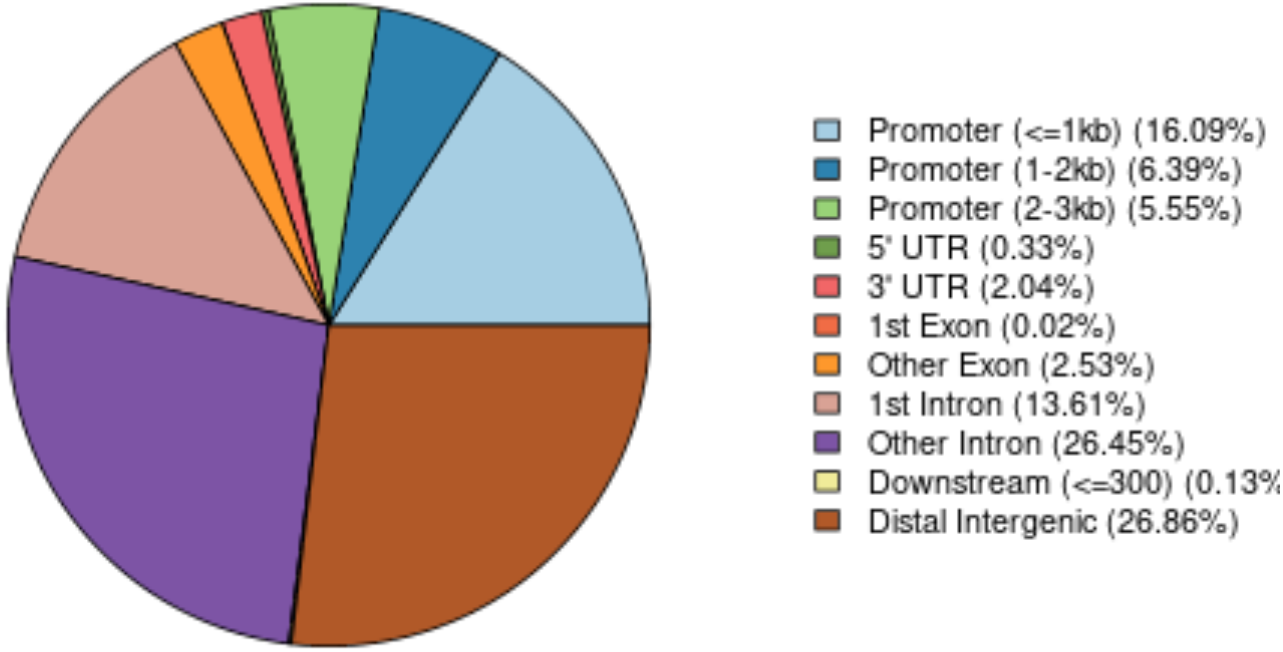
Visualization of genomic features annotation from. `ChIPSeeker`

**Figure S4:**
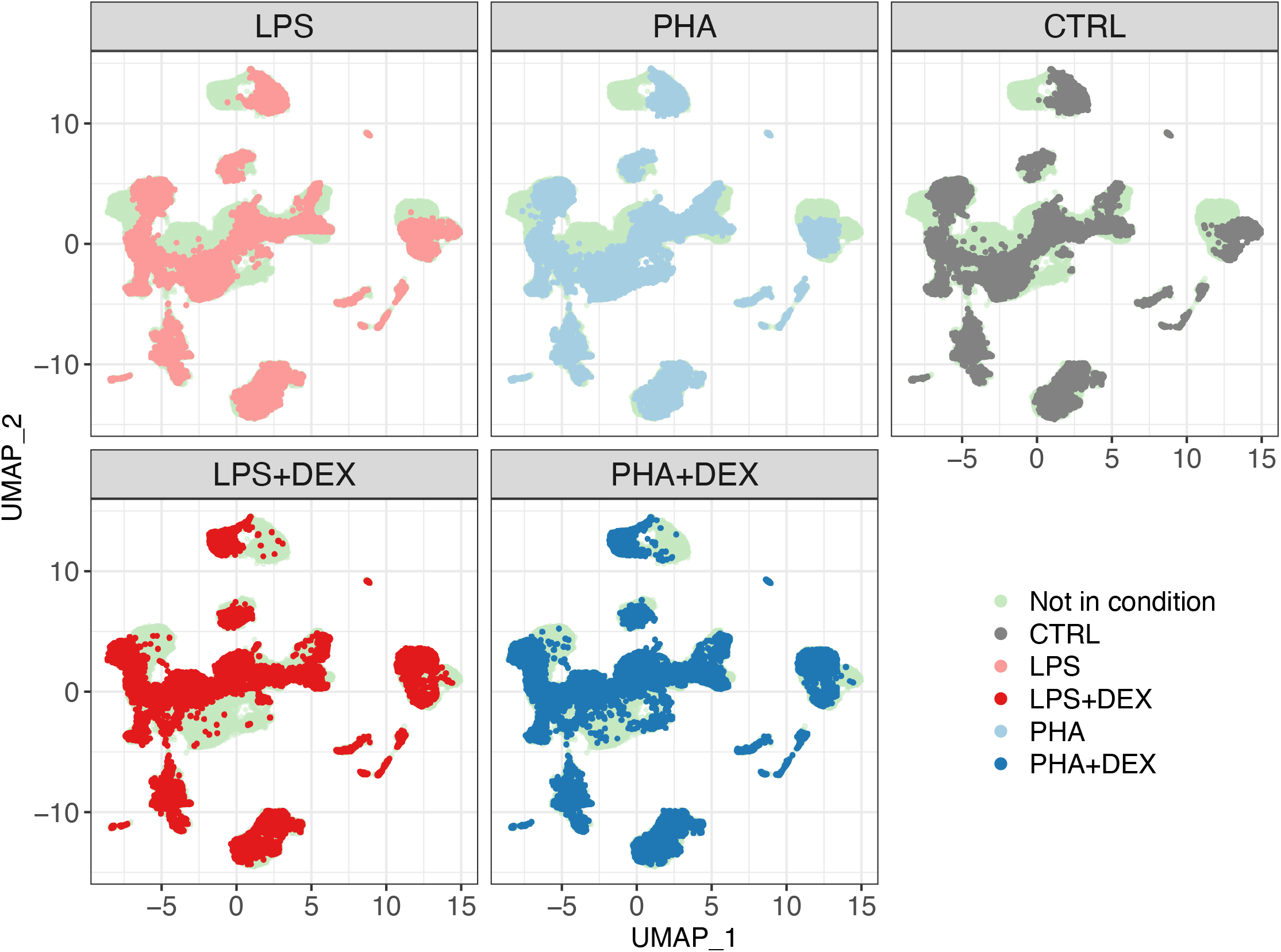
UMAP visualization of scATAC-seq data for each immune condition separately. Cells from each specific treatment condition are colored in grey (control), light red (LPS), red (LPS+DEX), light blue (PHA), and blue (PHA+DEX). Underneath, cells not present in the given condition are colored in light green.

**Figure S5:**
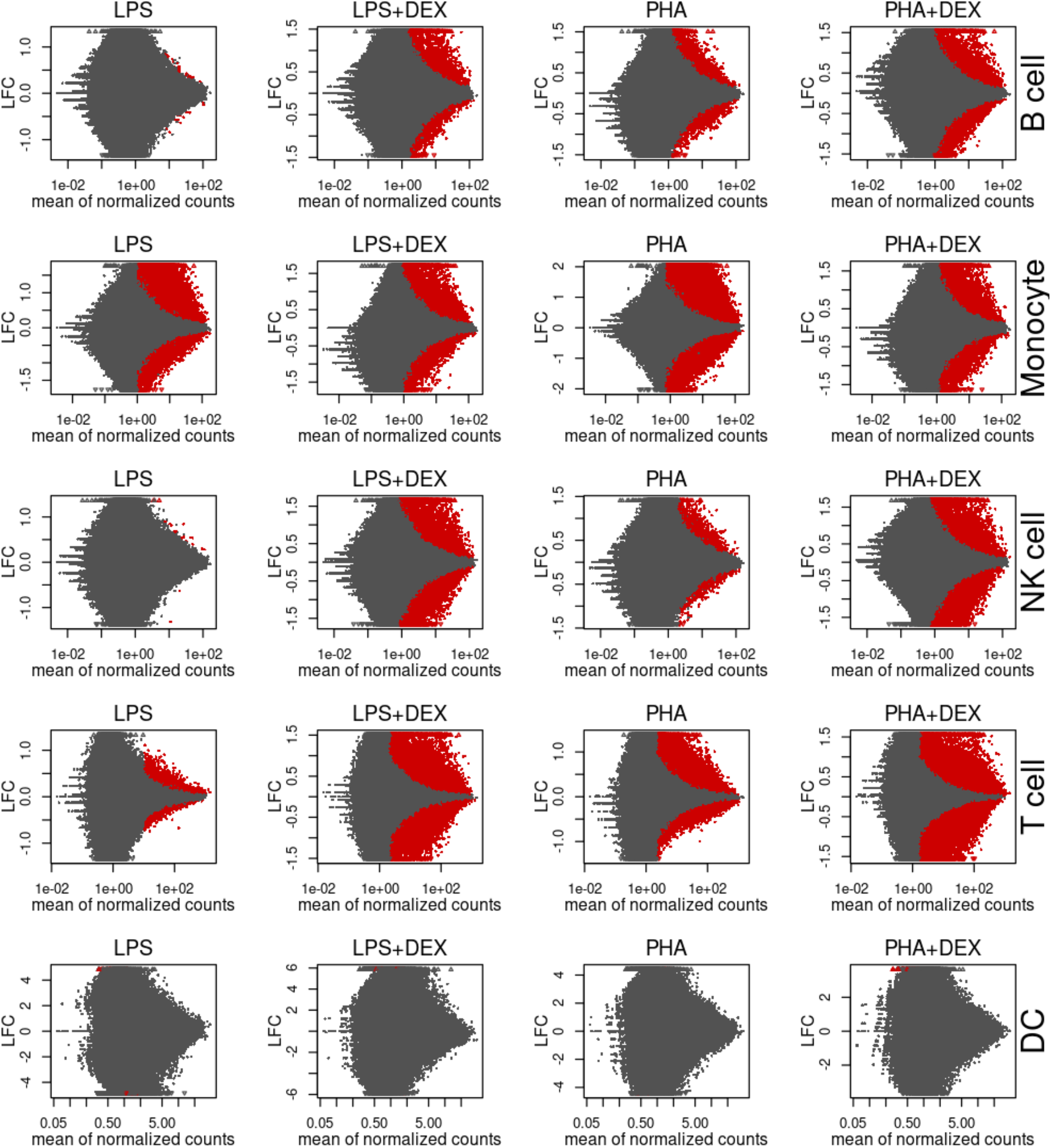
MA plots of differential chromatin accessibility analysis: rows representing five cell types (B cell, Monocyte, NK cell, T cell and DC) and columns representing 4 contrasts (LPS, LPS+DEX, PHA and PHA+DEX), for each panel the scatterplots of the log2 fold change (y-axis) of each peak against its mean of normalized counts (x-axis), which is calculated by the average of the normalized count values across *m* samples 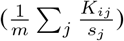. Here *K*_*ij*_ denotes read counts for *i*_*th*_ gene in *j*_*th*_ sample and *s*_*j*_ denoting the size factor to adjust the difference in sequence depth across samples, calculated from median-of-ratios method. And red color denoting peaks with FDR*<*0.1.

**Figure S6:**
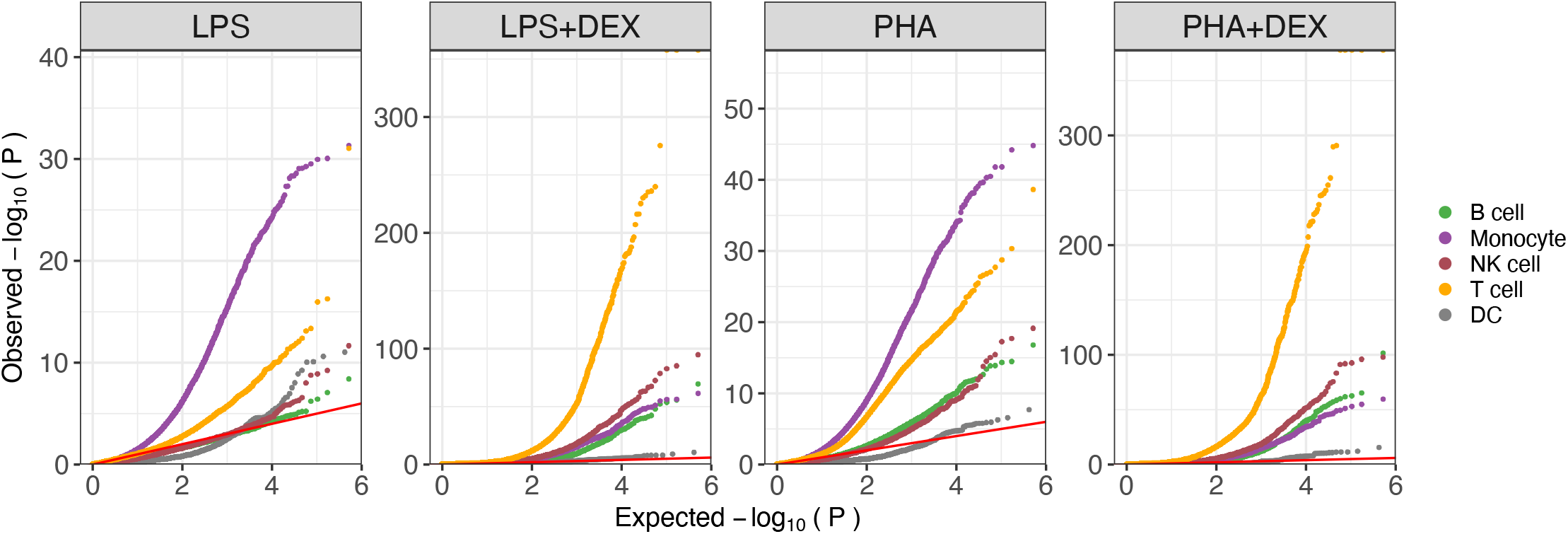
QQ plots of differential chromatin accessibility analysis colored by cell types for each contrast.

**Figure S7:**
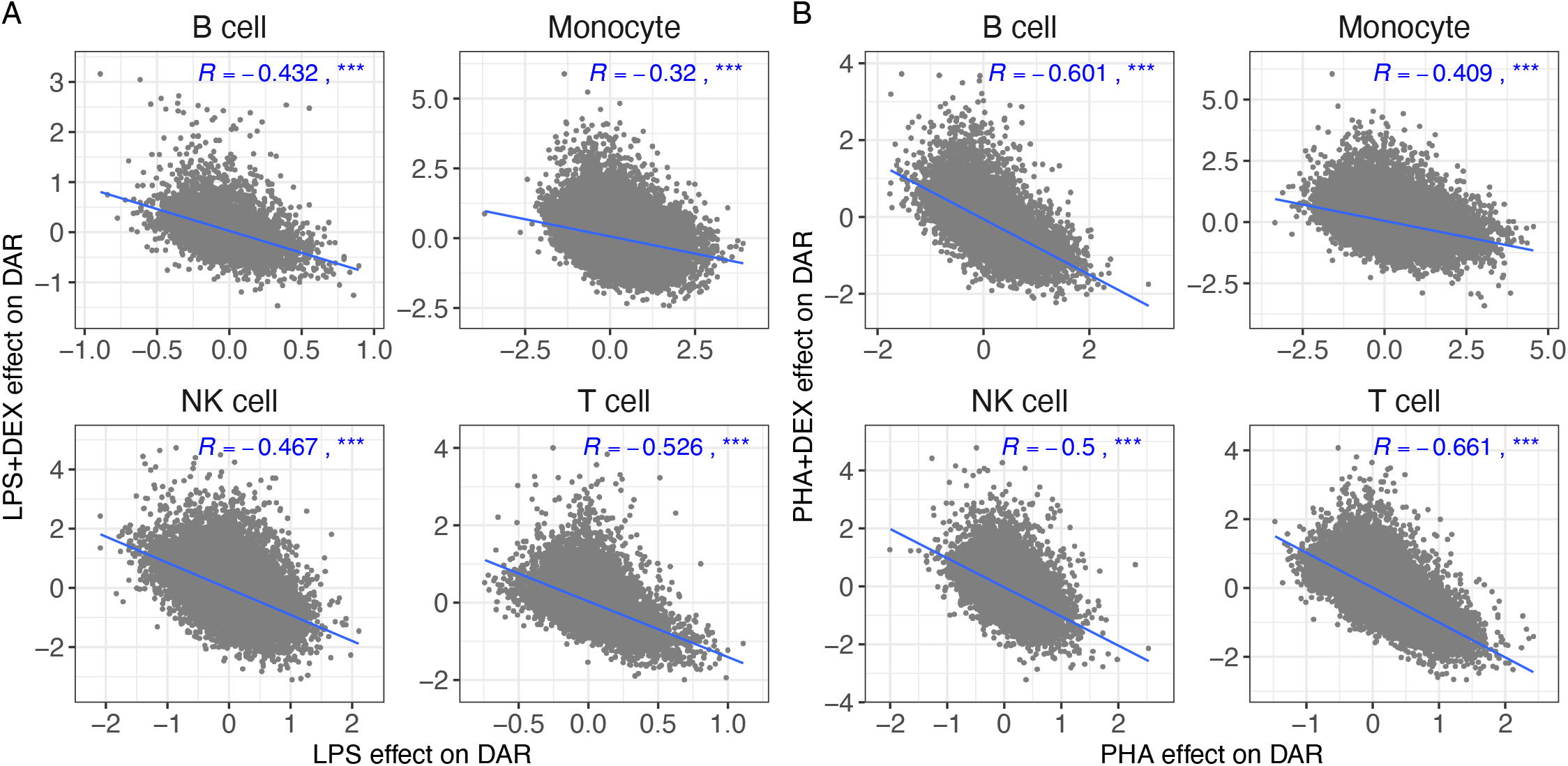
Dexamethasone (DEX) reverses effects of activation by immune stimuli on chromatin accessibility:. **(A)** Scatterplots of DEX effect on chromatin accessibility (y axis) against LPS effect on chromatin accessibility (x axis) **(B)** Scatterplots of DEX effect on chromatin accessibility (y axis) against PHA effect on chromatin accessibility (x axis). Blue lines represent the linear trend between the effects of immune stimuli and DEX treatments on chromatin accessibility. *R* represents the Pearson correlation coefficients of the effects of two types of immune treatments on chromatin accessibility, *denoting *p <* 0.05, ** denoting *p <* 0.01 and *** denoting *p*< 0.001.

**Figure S8:**
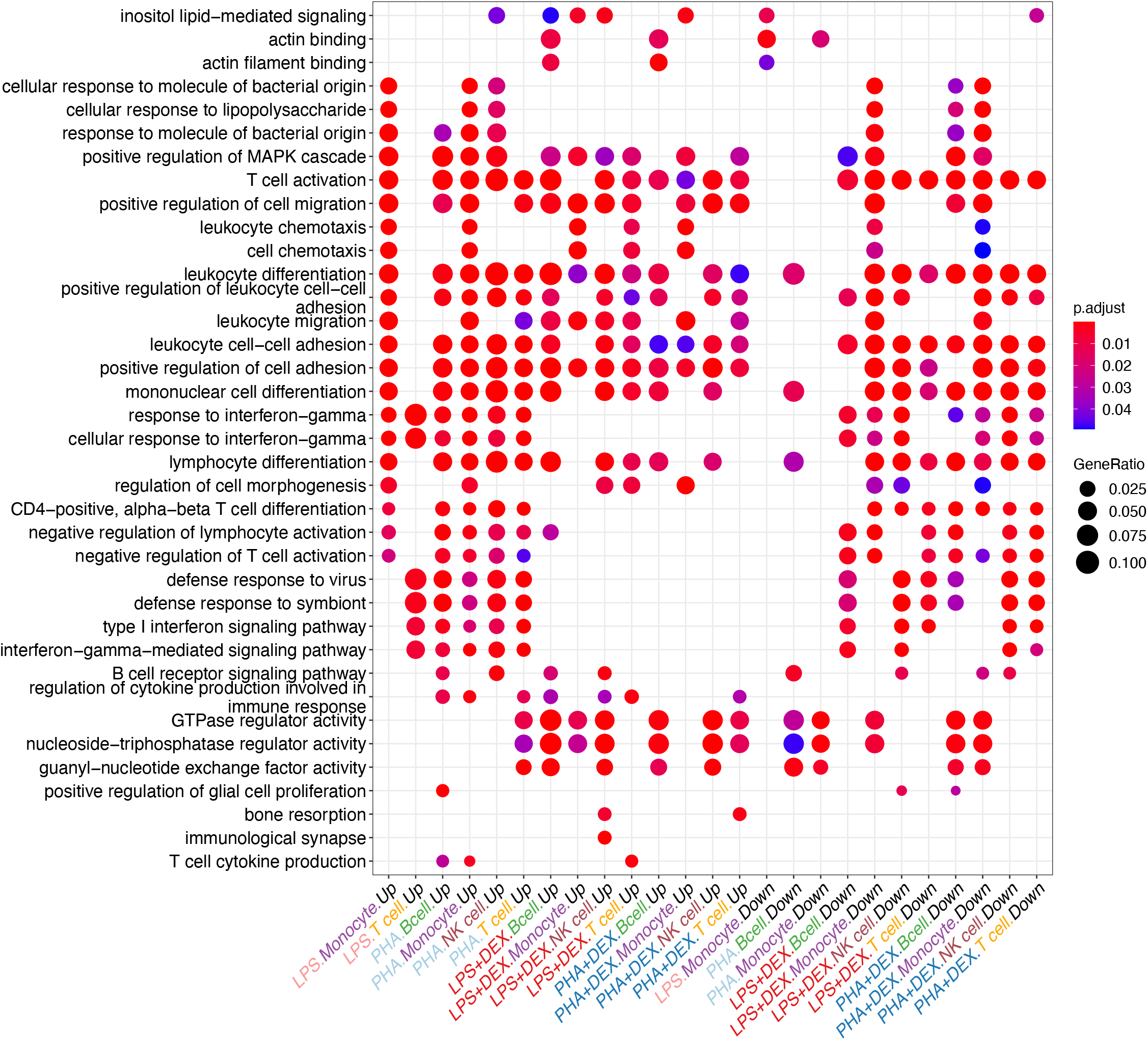
Visualization of Gene Ontology (GO) enrichment analysis for the closest genes in the DARs.

**Figure S9:**
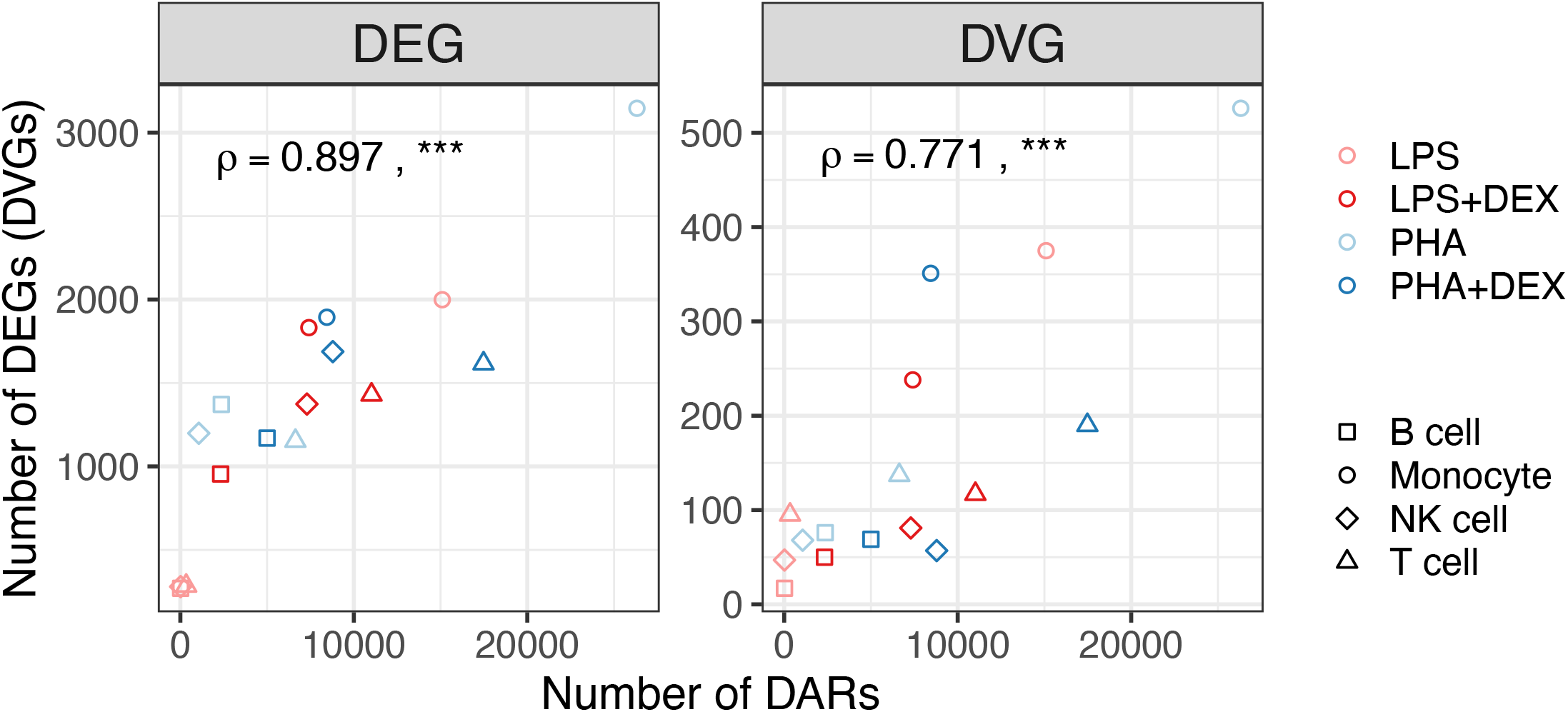
**Scatterplots of Number of DEGs (left panel) and DVGs (right panel) against number of DARs in corresponding conditions**, *** denoting a significant Spearman’s correlation with *p*< 0.001.

**Figure S10:**
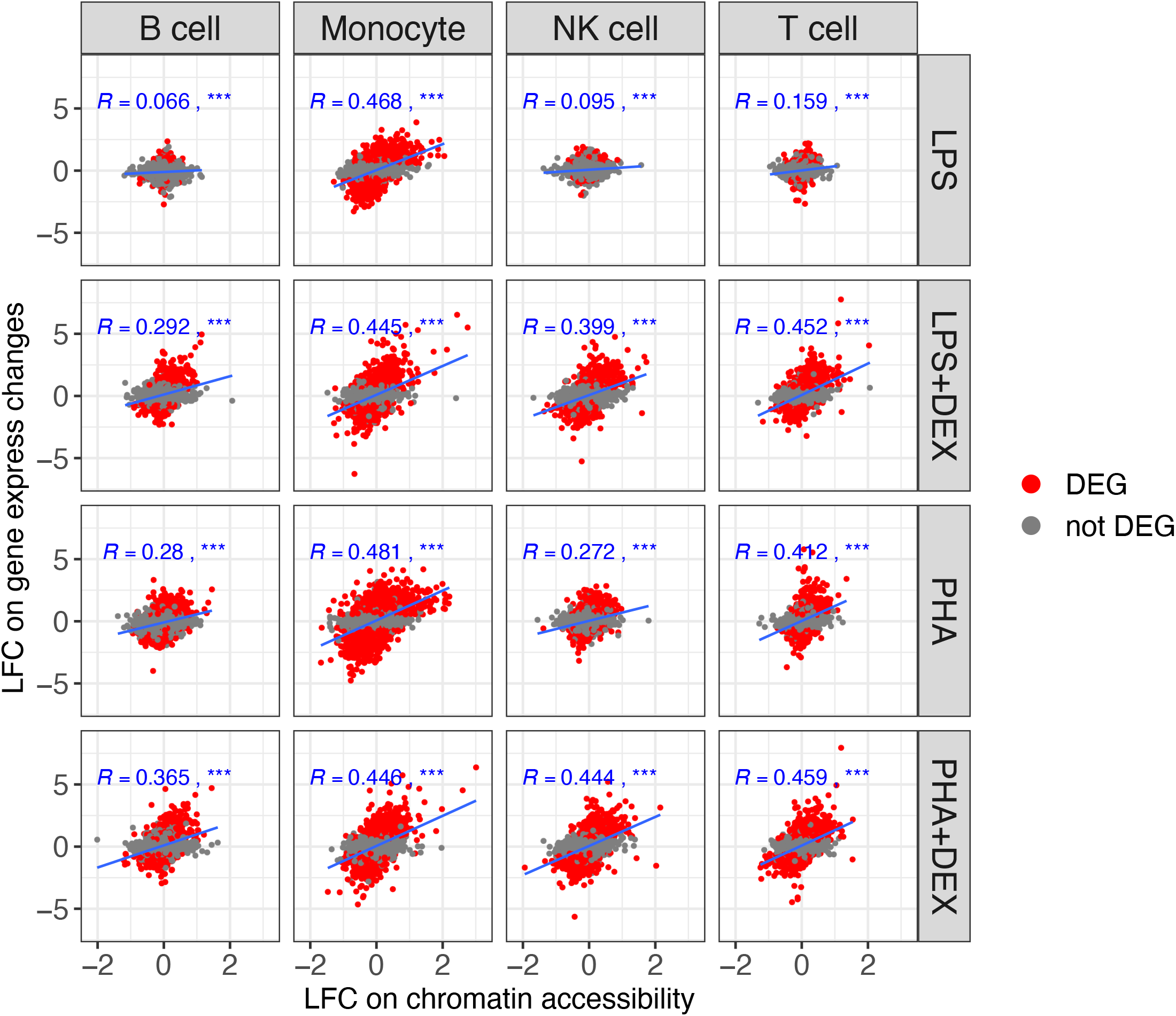
Scatterplots of LFC on gene expression (y-axis) against LFC on gene chromatin accessibility (x-axis). Blue lines represent the linear trend between response changes in gene expression and gene chromatin accessibility, *R* represents the Pearson correlation coefficients between response changes in the two types of molecular profiles, *** denoting a significant correlation(*p <* 0.001).

**Figure S11:**
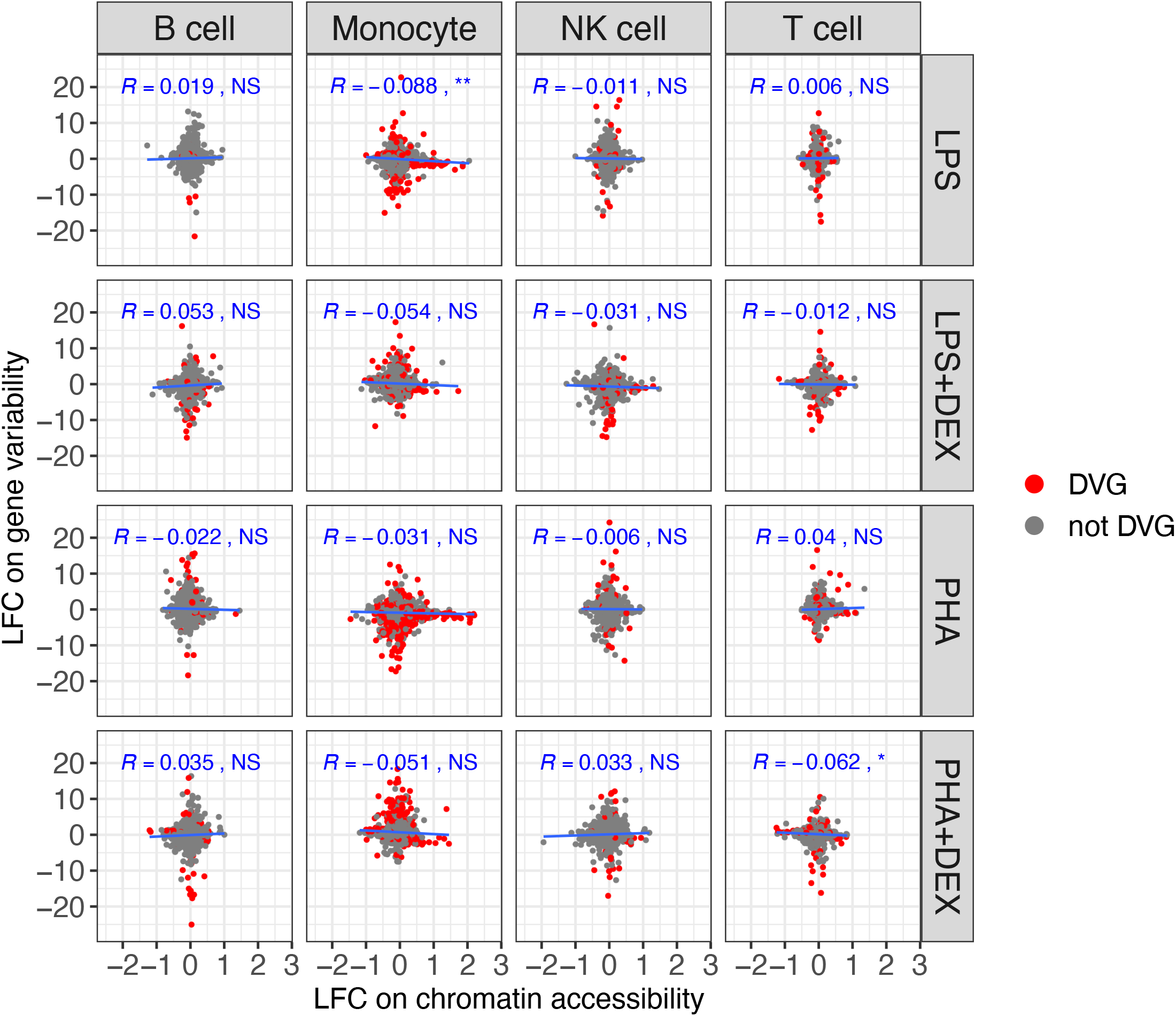
Scatterplots of LFC on gene variability (y-axis) against LFC on gene chromatin accessibility (x-axis). Blue lines represent the linear trend between response changes in gene variability and gene chromatin accessibility, *R* represents the Pearson correlation coefficients between response changes in the two types of molecular profiles, NS denoting not significance with *p>* 0.05, * denoting *p*< 0.05, ** denoting *p*< 0.01 and *** denoting *p*< 0.001.

**Figure S12:**
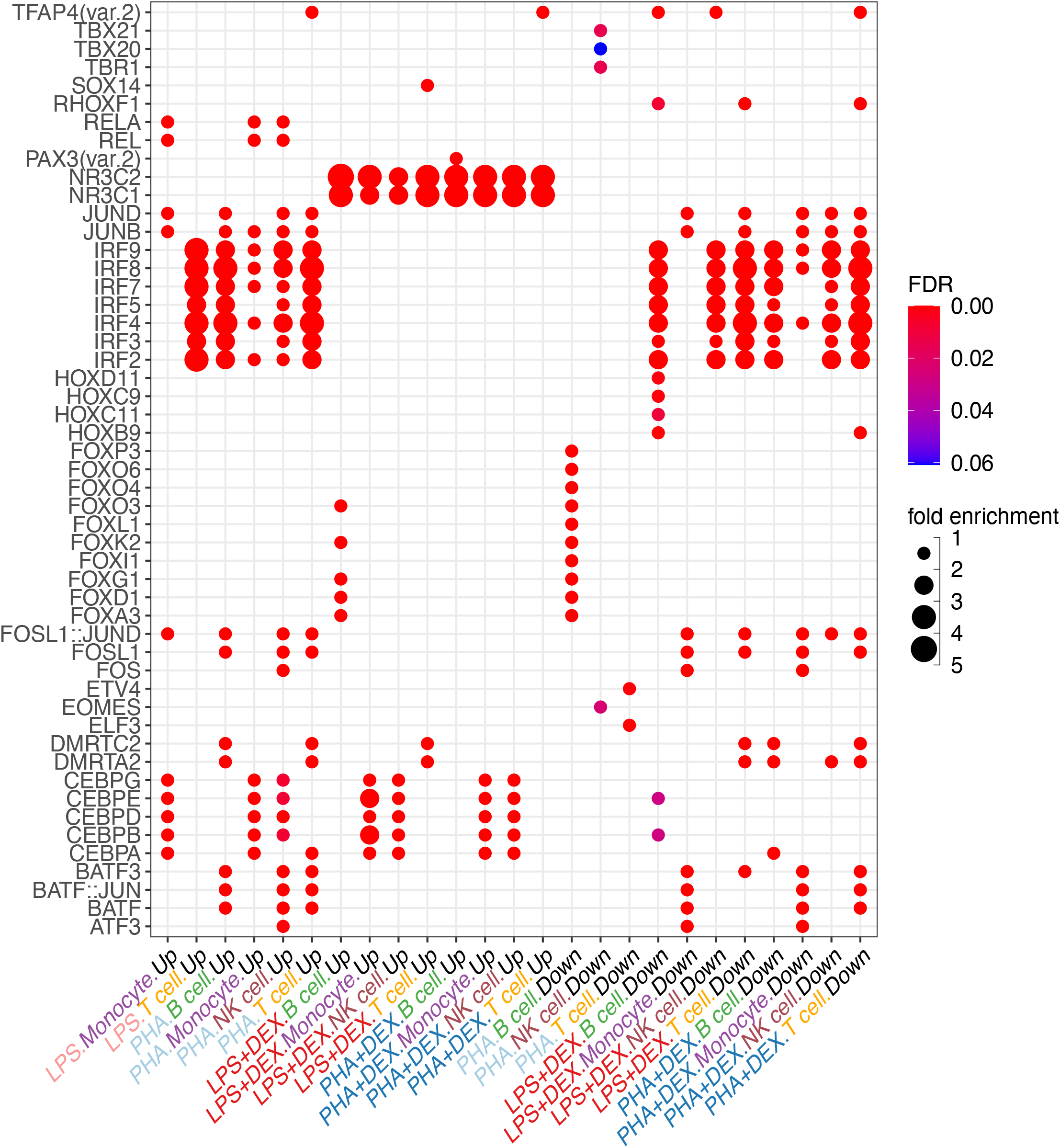
Visualization of enrichment of TF motifs in DARs: each dot represents a top enriched TF motif in the DARs for each condition (cell type+contrast) and direction (increased or decreased).

**Figure S13:**
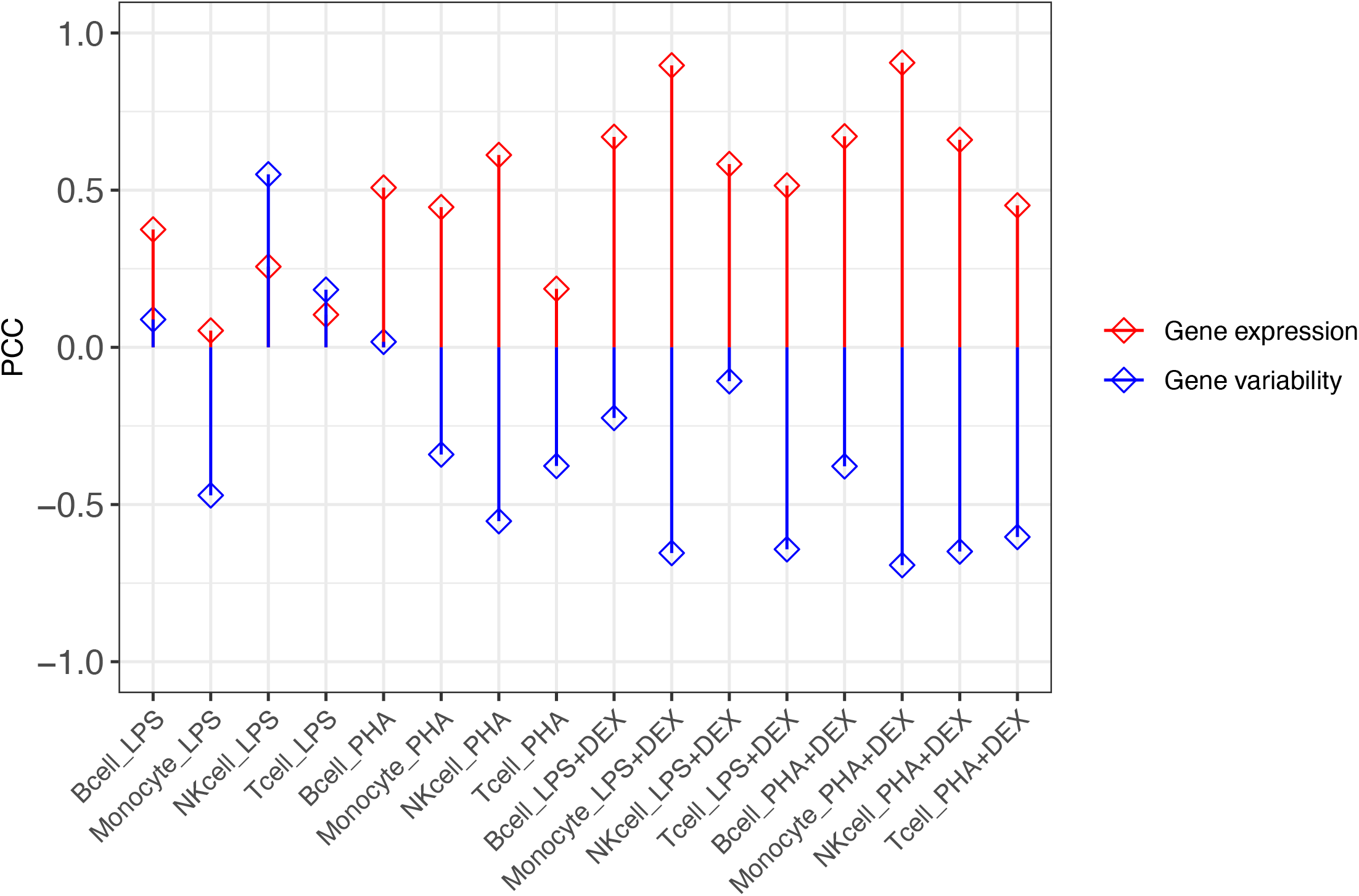
Pearson correlation coefficient (PCC) in 16 conditions (4 cell type × 4 contrast) Between changes on TF regulated gene expression (red color) or gene variability (blue color) with changes on TF activities.

**Figure S14:**
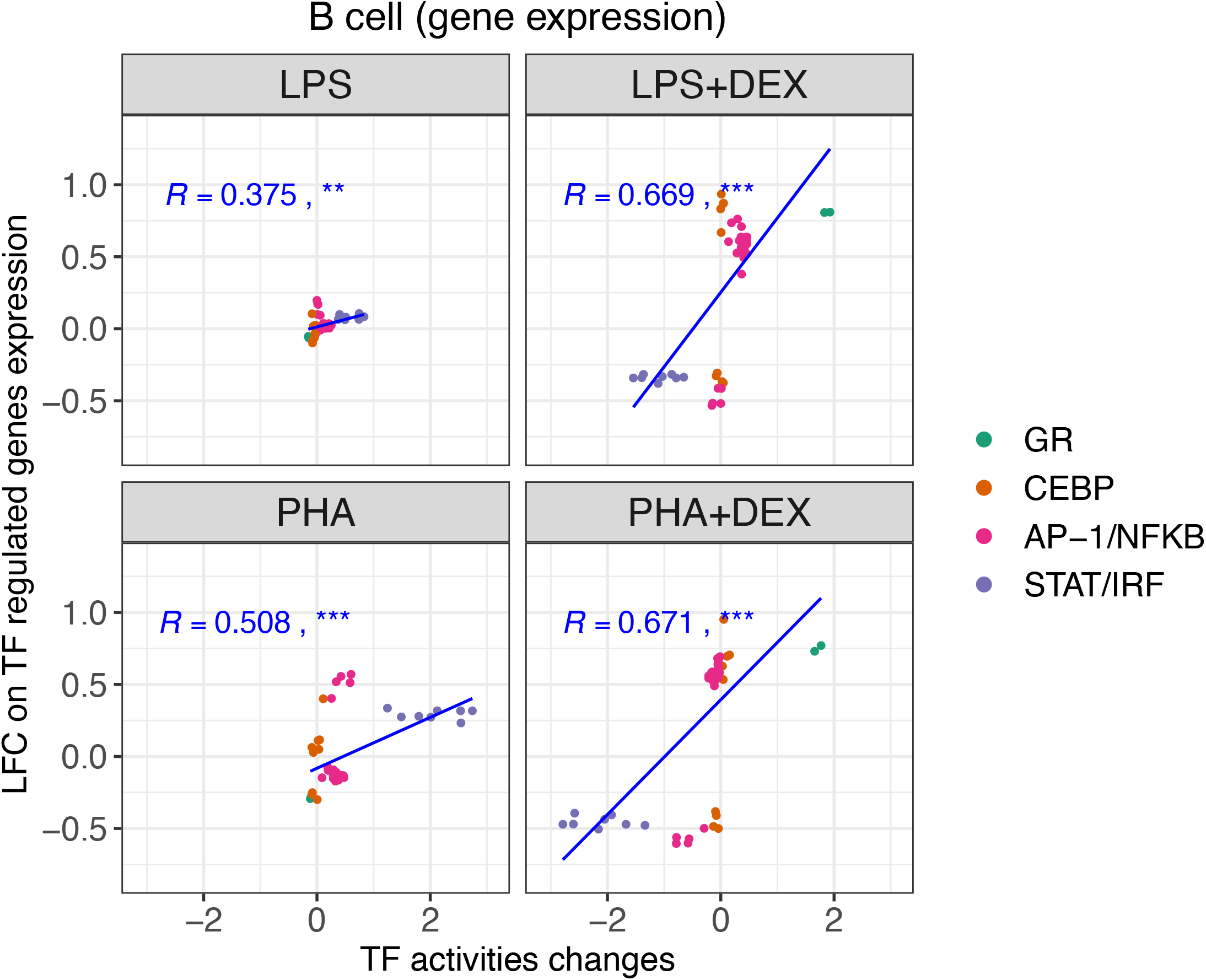
**Scatterplots of Changes of TF motif regulated gene expression against those of TF motif activities in B cells**, each panel representing LPS, LPS+DEX, PHA and PHA+DEX, respectively and colored by 4 different patterns of TF motifs. R value represents Pearson correlation coefficient, NS not significant, * *p <* 0.05, ** *p <* 0.01 and *** *p <* 0.001. Blue lines represent the regression between changes of TF motif regulated gene expression against changes of TF motif activities.

**Figure S15:**
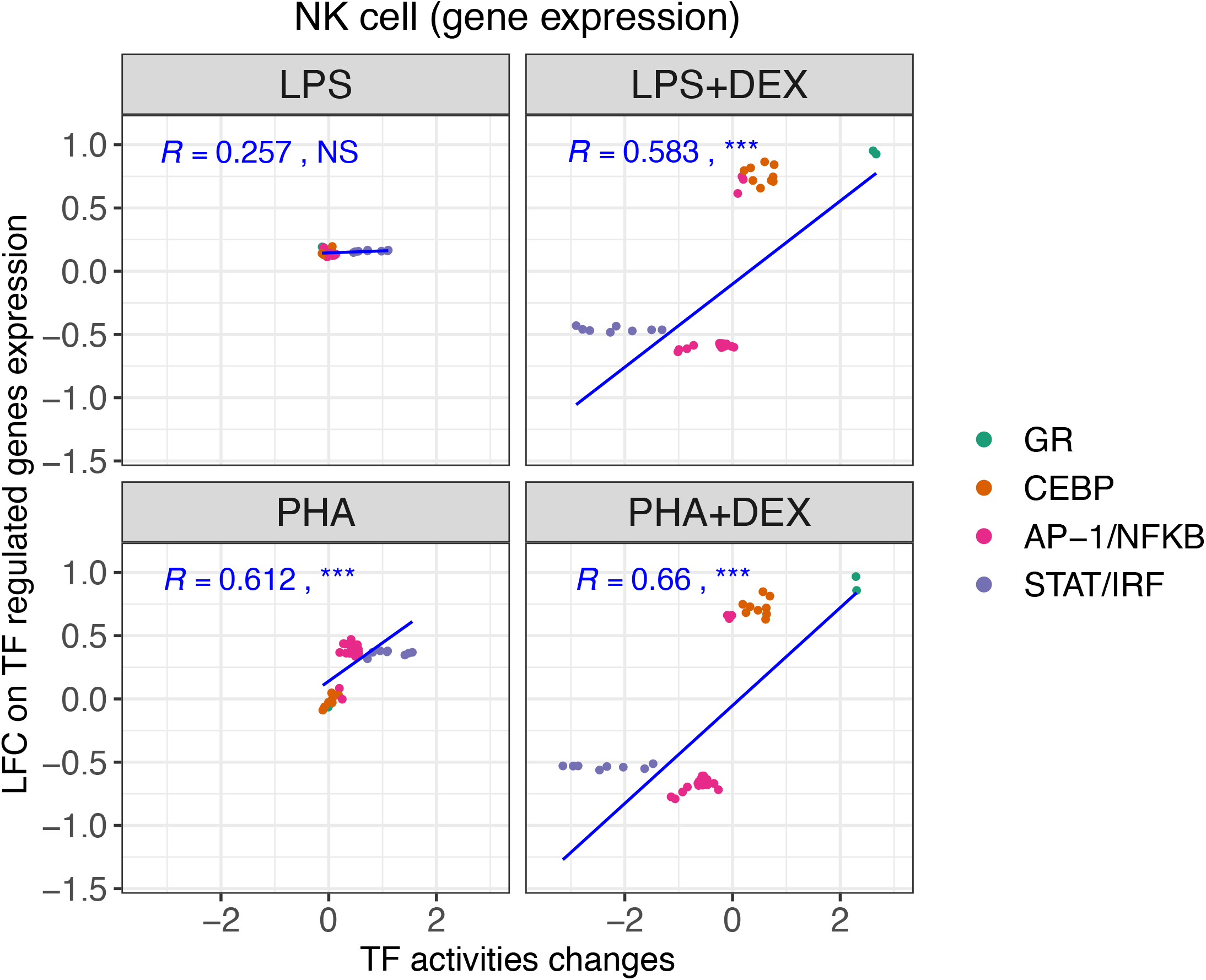
Scatterplots of Changes of TF motif regulated gene expression against those of TF motif activities in NK cells, each dot for TF motif and colored by 4 different patterns of TF motif. Each panel represents LPS, LPS+DEX, PHA and PHA+DEX, respectively. R value represents Pearson correlation coefficient, NS not significant, * *p*< 0.05, ** *p*< 0.01 and *** *p*< 0.001. Blue lines represent the regression between changes of TF motif regulated gene expression against changes of TF motif activities.

**Figure S16:**
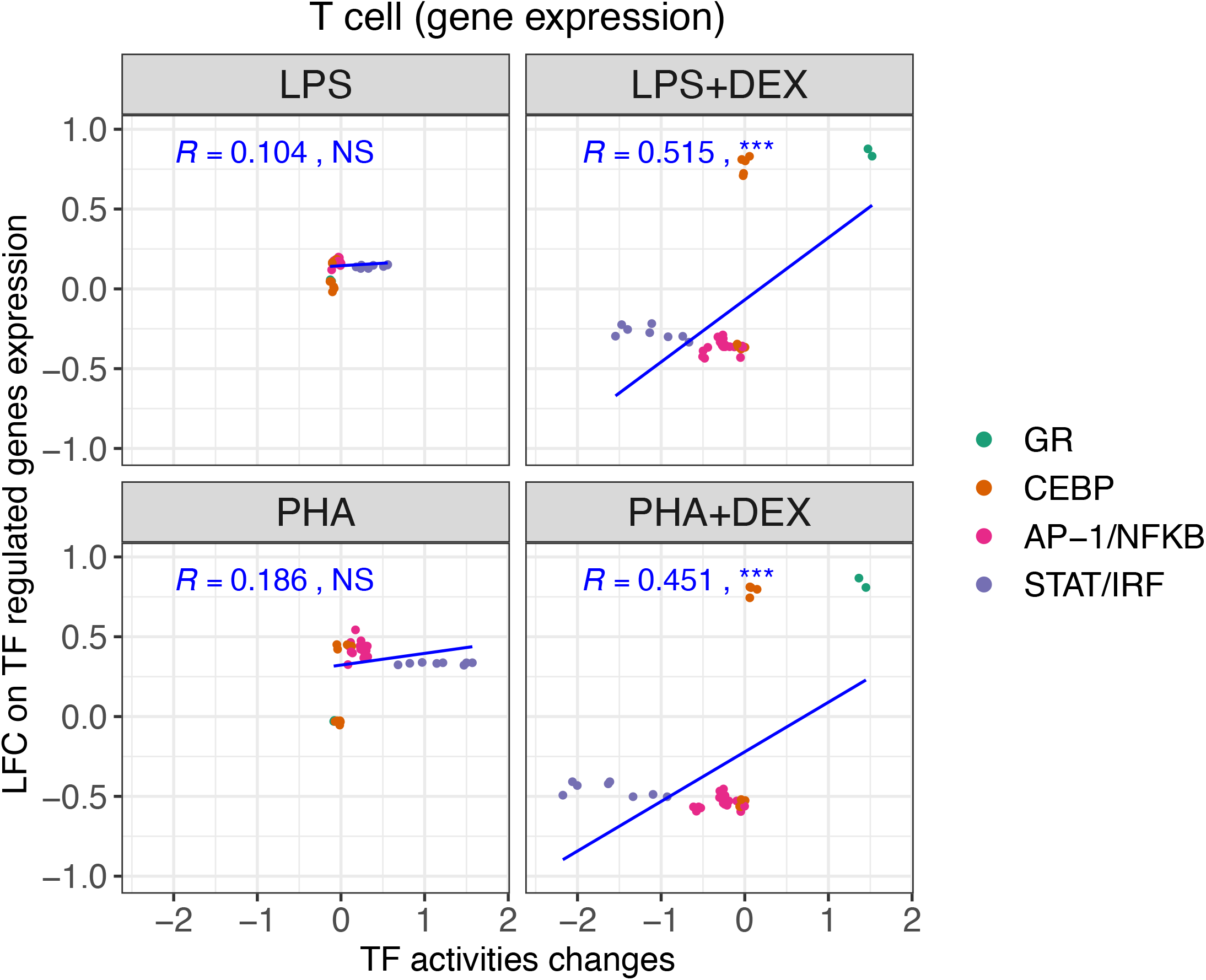
Scatterplots of Changes of TF motif regulated gene expression against those of TF motif activities in T cells, each dot for TF motif and colored by 4 different patterns of TF motif. Each panel represents LPS, LPS+DEX, PHA and PHA+DEX, respectively. R value represents Pearson correlation coefficient, NS not significant, * *p*< 0.05, ** *p*< 0.01 and *** *p*< 0.001. Blue lines represent the regression between changes of TF motif regulated gene expression against changes of TF motif activities.

**Figure S17:**
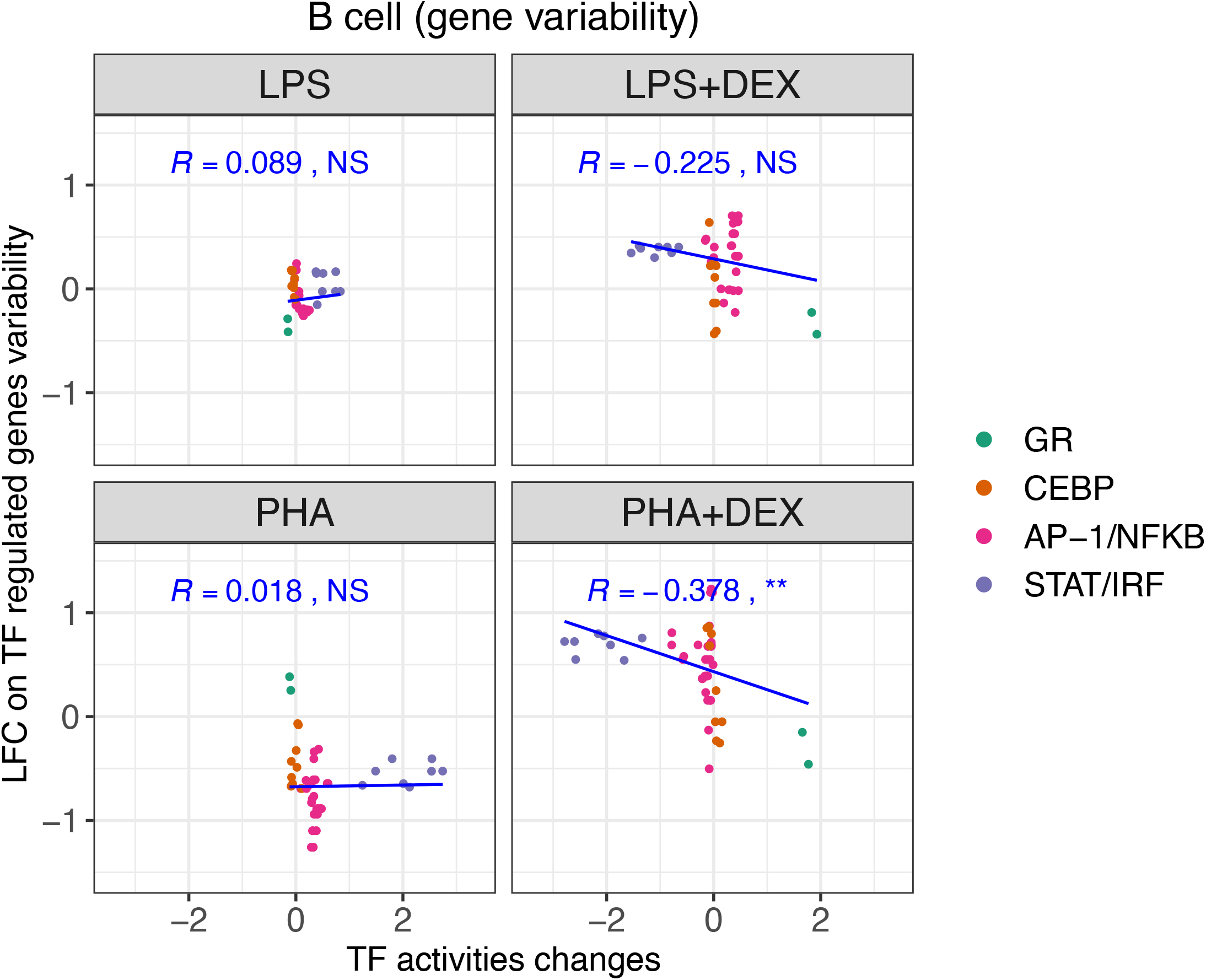
Scatterplots of Changes of TF motif regulated gene variability against those of TF motif activities in B cells, each panel representing LPS, LPS+DEX, PHA and PHA+DEX, respectively and colored by 4 different patterns of TF motifs. R value represents Pearson correlation coefficient, NS not significant, * *p <* 0.05, ** *p <* 0.01 and *** *p <* 0.001. Blue lines represent the regression between changes of TF motif regulated gene variability against changes of TF motif activities.

**Figure S18:**
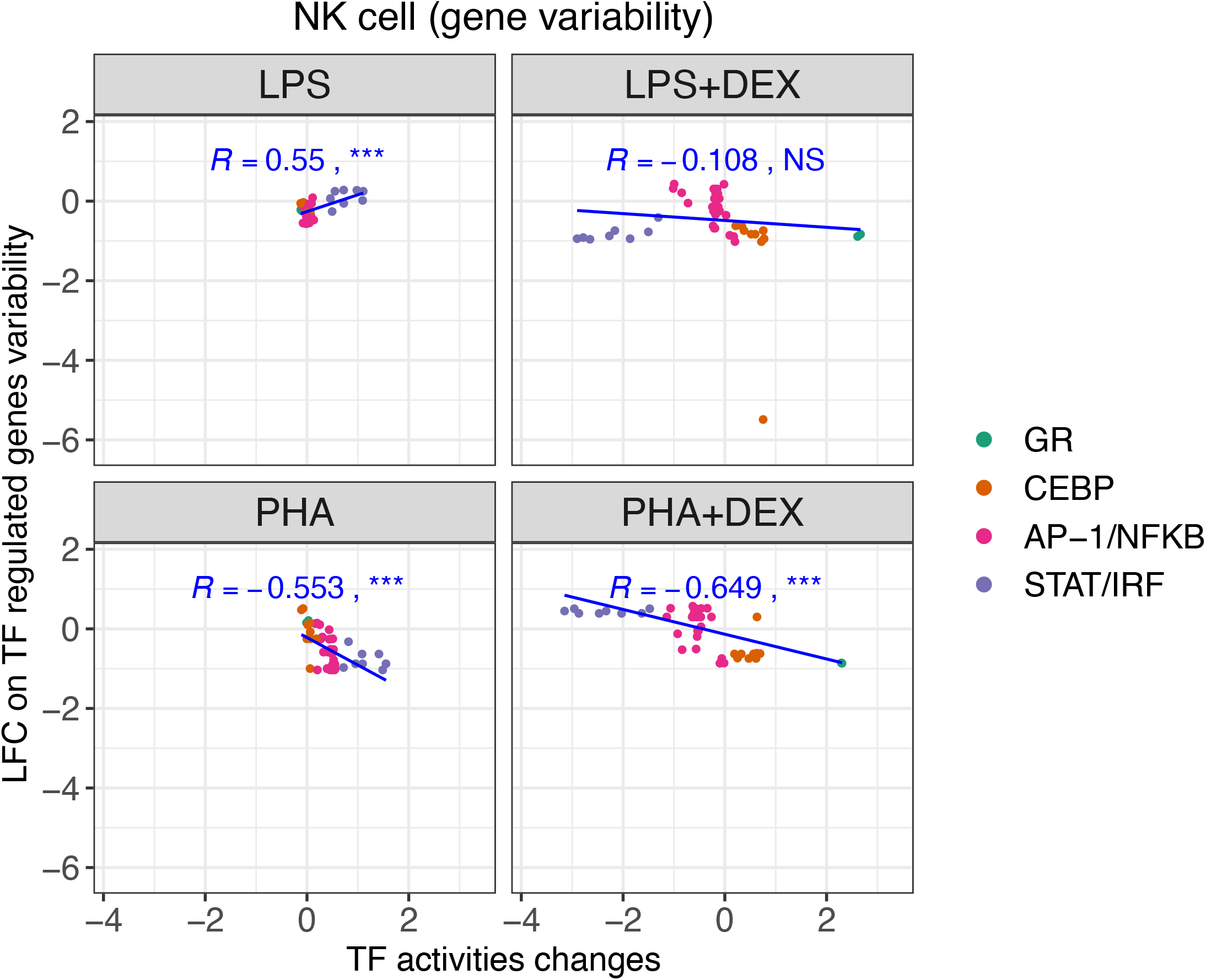
Scatterplots of Changes of TF motif regulated gene variability against those of TF motif activities in NK cells, each panel representing LPS, LPS+DEX, PHA and PHA+DEX, respectively and colored by 4 different patterns of TF motifs. R value represents Pearson correlation coefficient, NS not significant, * *p <* 0.05, ** *p <* 0.01 and *** *p <* 0.001. Blue lines represent the regression between changes of TF motif regulated gene variability against changes of TF motif activities.

**Figure S19:**
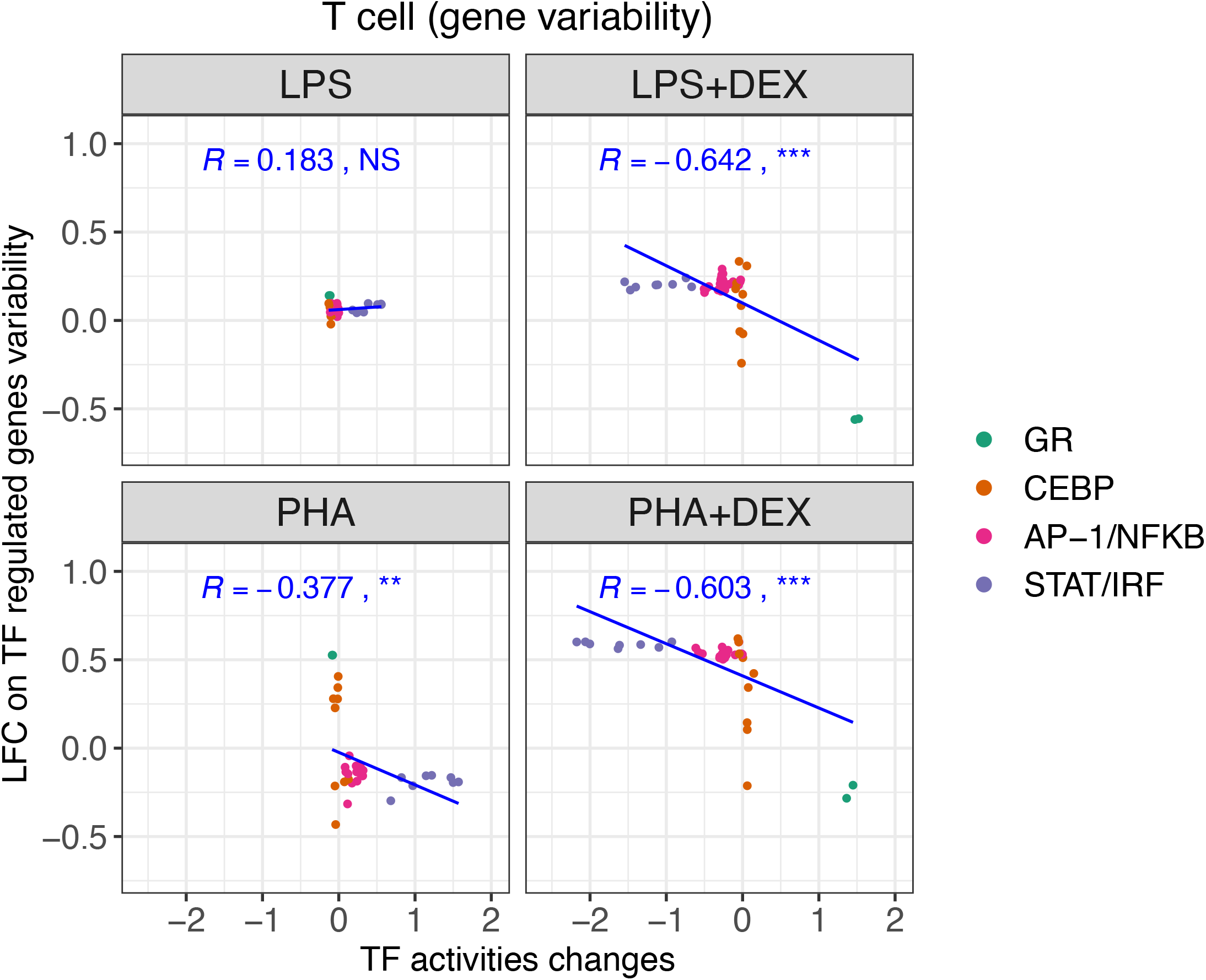
Scatterplots of Changes of TF motif regulated gene variability against those of TF motif activities in T cells, each panel representing LPS, LPS+DEX, PHA and PHA+DEX, respectively and colored by 4 different patterns of TF motifs. R value represents Pearson correlation coefficient, NS not significant, * *p <* 0.05, ** *p <* 0.01 and *** *p <* 0.001. Blue lines represent the regression between changes of TF motif regulated gene variability against changes of TF motif activities..

**Figure S20:**
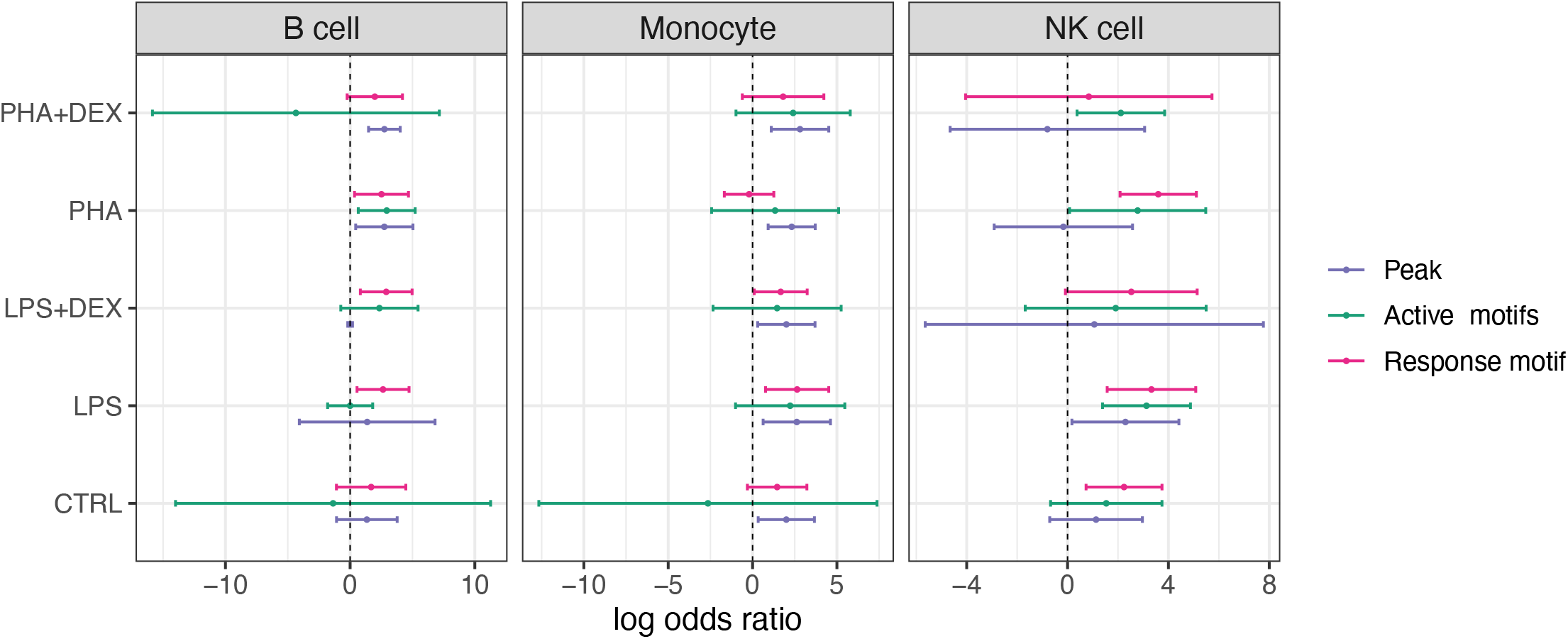
Enrichment of eQTL signals in three genomic features in single-cell eQTL mapping datasets, pink denoting variants in Response motif, green denoting variants in Active motifs and blue for the remaining variants in the peak.

**Figure S21:**
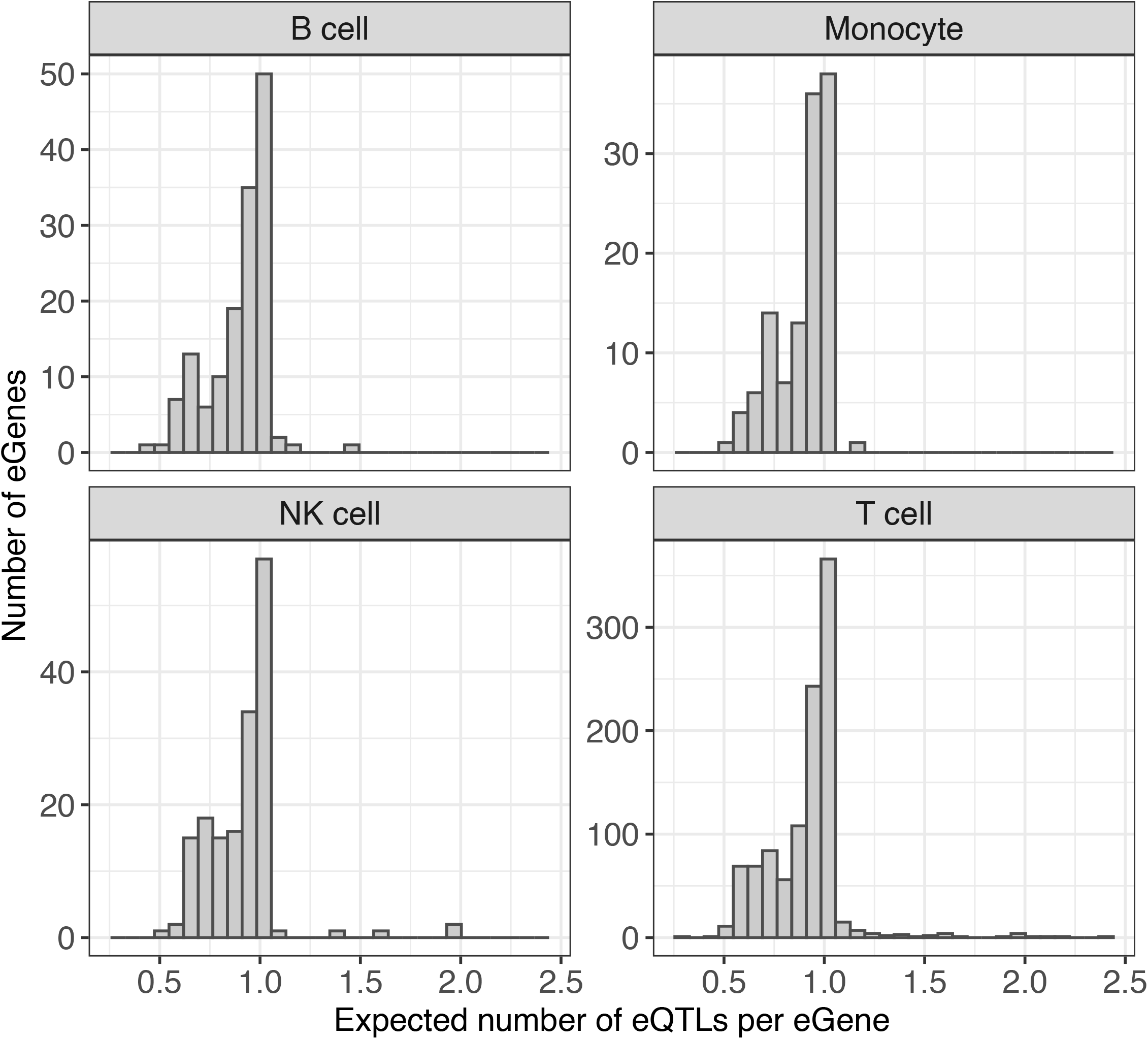
Expected number of eQTLs per eGene across conditions in B cell, Monocyte, NK cell and T cell.

**Figure S22:**
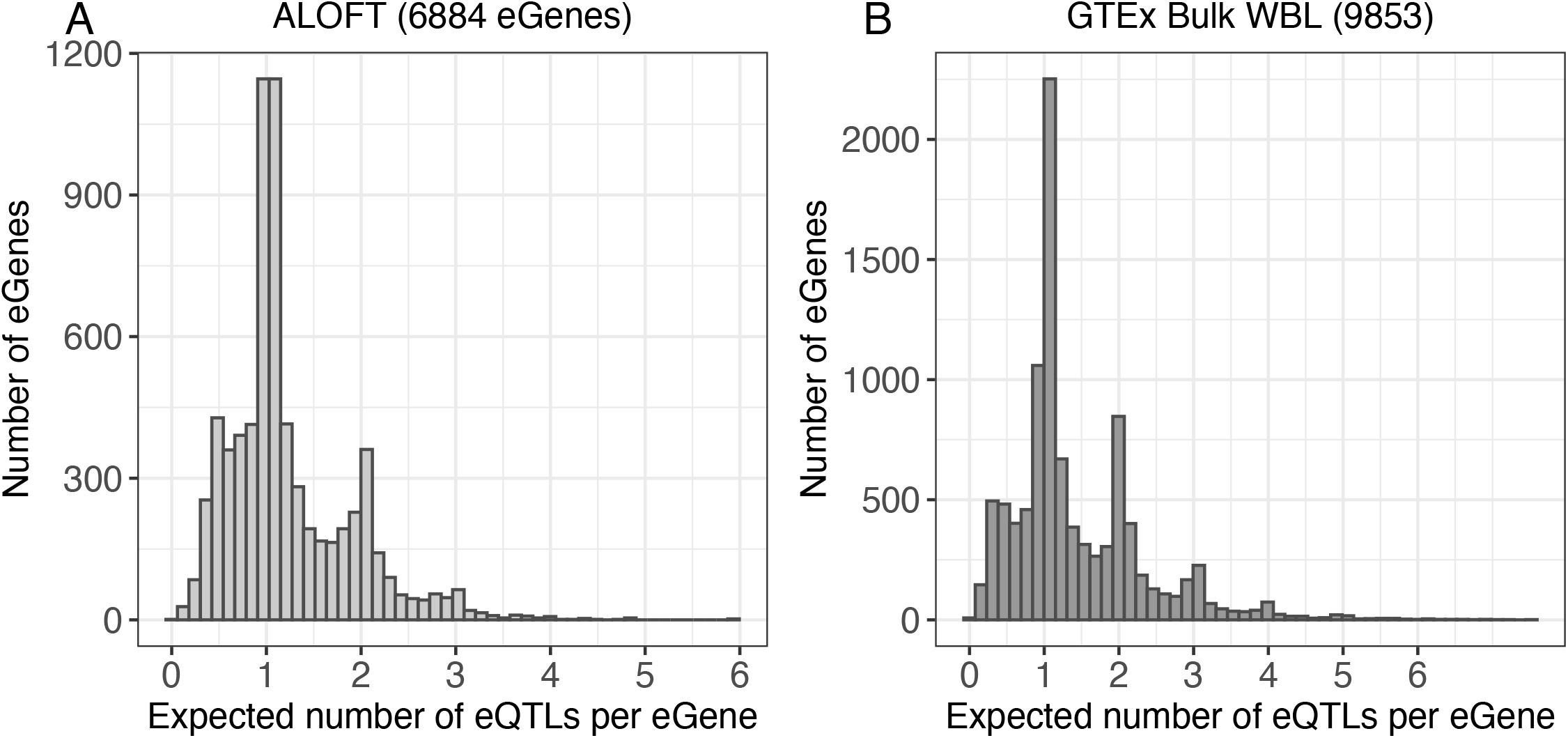
Expected number of eQTLs per eGene detected in ALOFT cohort (A) and GTEx whole blood (B).

**Figure S23:**
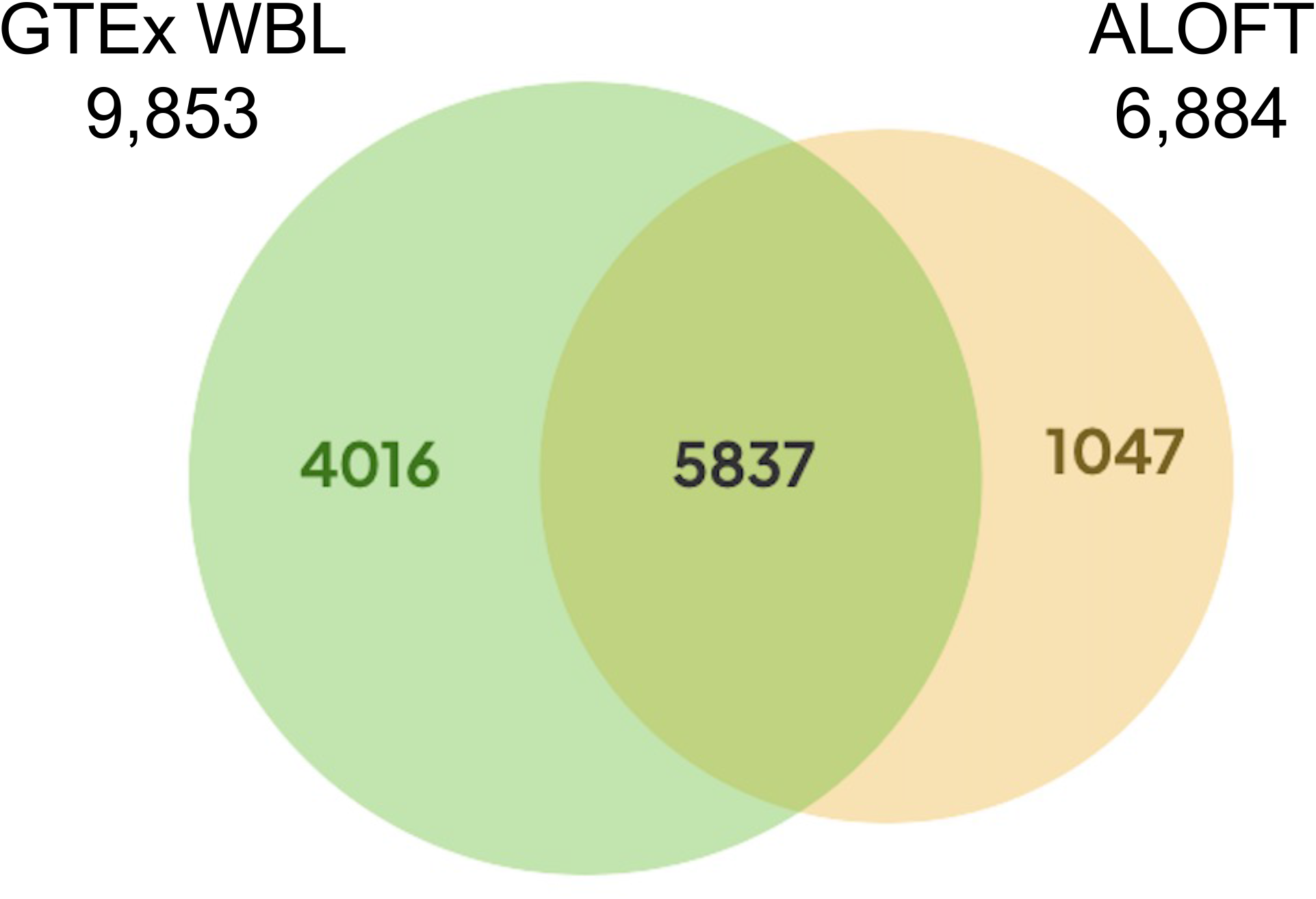
Venn diagram showing overlap eGenes identified in ALOFT cohort and GTEx whole blood.

**Figure S24:**
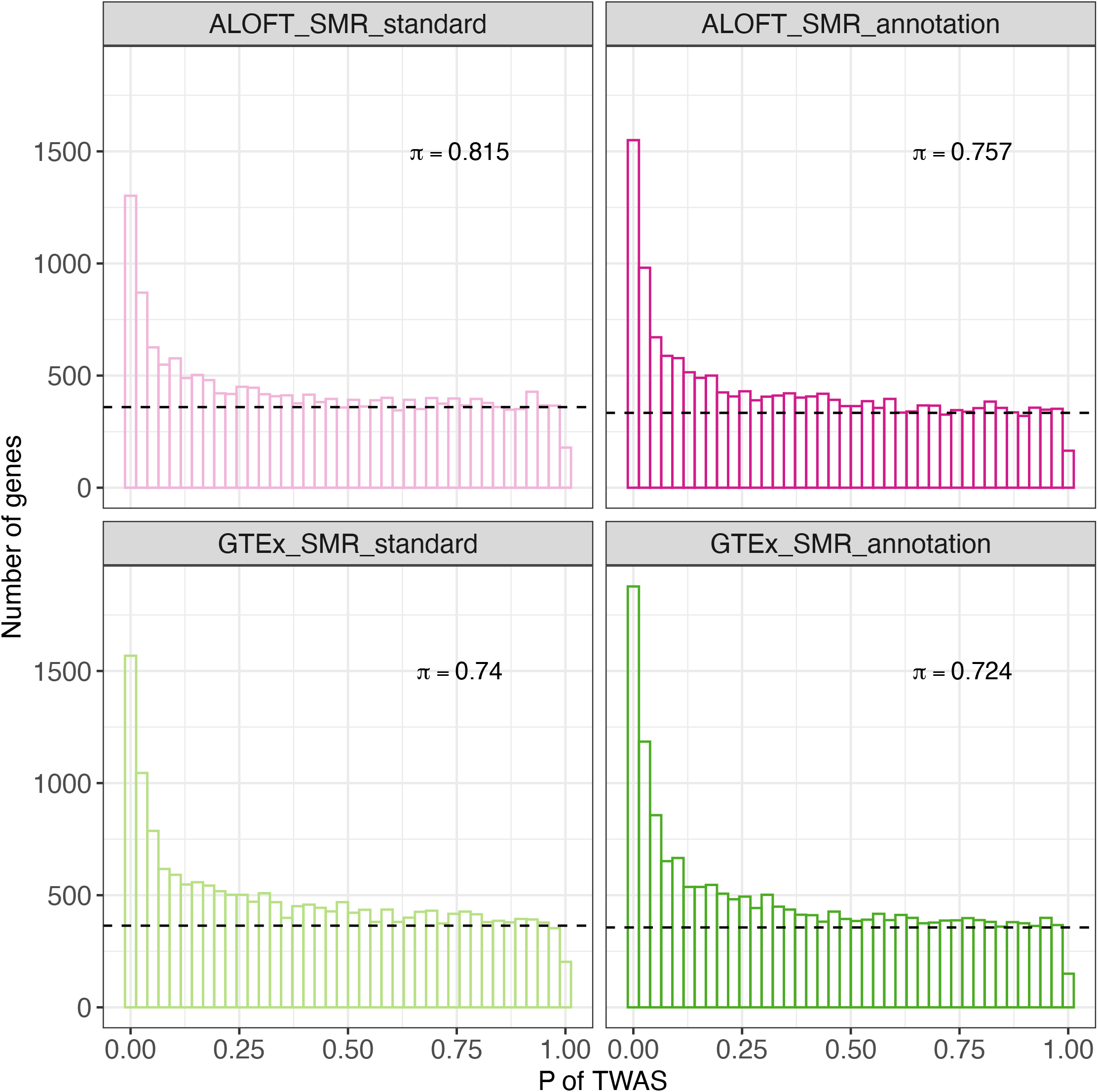
Histogram of distribution of *p*-value from the TWAS/SMR analysis in the ALOFT cohort and GTEx whole blood using two methods. (left) using minimum *p*-value or (right) the maximum PIP from fine-mapping eQTLs from DAP-G. *π* is the proportion of true null associated genes estimated from the `qvalue` package and the dashed line is the expected number of genes under the null hypothesis for each bin.

**Figure S25:**
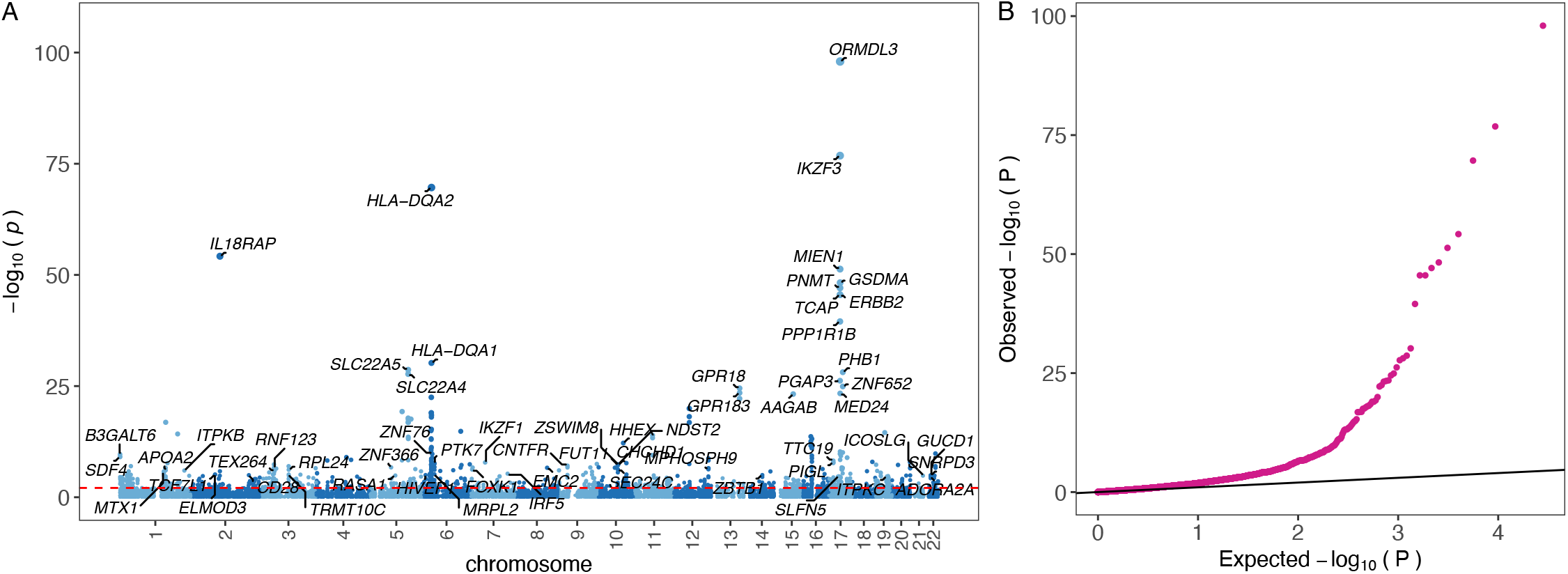
TWAS of asthma using GTEx cohort from whole blood tissue. **A** Manhattan plot visualization of genes genetically associated with asthma risk with TWAS with the annotation from GTEx cohort, each dot representing the SMR *-* log_10_(*p*) of each gene in the y-axis. **B** Quantile-quantile (QQ) plot for TWAS with the annotation from the GTEx cohort.

**Figure S26:**
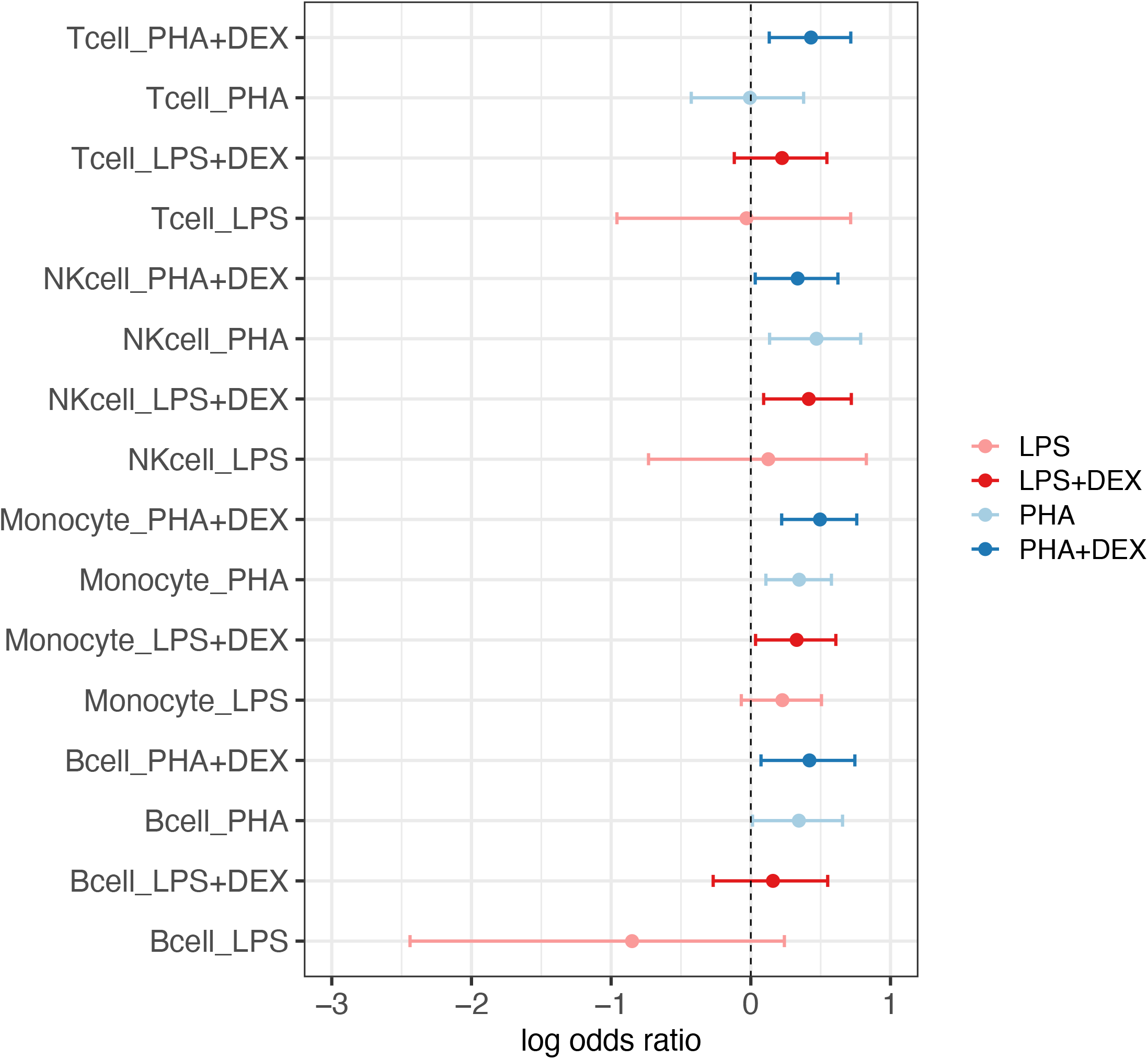
Forest plot of the log odds ratio of asthma-risk genes enriched in DEGs that were identified across 16 conditions.

## Supplementary Tables

**Table S1:** Demographic table of 96 individuals in the scRNA-seq experiment. SD= Standard Deviation. FFPP= FEV1/FVC percent predicted (FFPP) is the ratio of maximum air expelled during the first second following maximal inhalation (FEV1) to the amount of air forcibly exhaled after maximal inhalation (FVC), converted to percentage of predicted based on patient demographics.

|  | No | Mean | Median | SD |
| --- | --- | --- | --- | --- |
| Sex |  |  |  |  |
| Male | 49 |  |  |  |
| Female | 47 |  |  |  |
| Age |  |  |  |  |
| 10.58-15.87 | 96 | 13.11 | 12.96 | 1.38 |
| FFPP |  |  |  |  |
| 26-113 | 91 | 86.01 | 88.00 | 14.58 |

**Table S2:** Demographic table of 16 individuals in the scATAC-seq experiment. SD= Standard Deviation. FFPP=FEV1/FVC percent predicted (FFPP) is the ratio of maximum air expelled during the first second following maximal inhalation (FEV1) to the amount of air forcibly exhaled after maximal inhalation (FVC), converted to percentage of predicted based on patient demographics.

|  | No | Mean | Median | SD |
| --- | --- | --- | --- | --- |
| Sex |  |  |  |  |
| Male | 8 |  |  |  |
| Female | 8 |  |  |  |
| Age |  |  |  |  |
| 10.84 - 14.23 | 16 | 12.40 | 12.52 | 0.93 |
| FFPP |  |  |  |  |
| 66-107 | 15 | 91.80 | 93.00 | 13.48 |

**Table S3:**
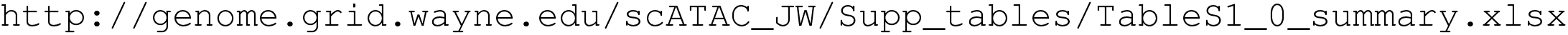
Summary metrics of cellranger-atac output. http://genome.grid.wayne.edu/scATAC_JW/Supp_tables/TableS1_0_summary.xlsx

**Table S4:** Summary statistics of single-cell data for each treatment. Column 2, number of individuals for each treatment; Column 3, median of number of cells per individual across individuals in each treatment; Column 4, median of average Reads per cell from the same individual across individuals in each treatment, Column 5, median of average features per cell from the same individual across individuals in each treatment.

|  | #Individuals | # Cells per individual (median) | #Reads per cell (median) | #Features per cell (median) |
| --- | --- | --- | --- | --- |
| CTRL | 16 | 1,280 | 2,588 | 5,997 |
| LPS | 16 | 1,030 | 2,451 | 6,738 |
| LPS+DEX | 16 | 1,118 | 2,672 | 7,240 |
| PHA | 16 | 1,339 | 2,981 | 6,495 |
| PHA+DEX | 16 | 1,394 | 2,618 | 5,886 |

**Table S5:** Summary of differential accessible regions for each condition. Column 2, total number of peaks in differential analysis; Column 3, number of differential accessible regions (DARs); Column 4, the proportion of DARs in the testing peaks.

| conditions | #testing peaks | #DAR | proportion |
| --- | --- | --- | --- |
| Bcell_LPS | 260111 | 18 | 0.007% |
| Bcell_LPS+DEX | 260111 | 2324 | 0.893% |
| Bcell_PHA | 260111 | 2363 | 0.908% |
| Bcell_PHA+DEX | 260111 | 5000 | 1.922% |
| Monocyte_LPS | 260756 | 15096 | 5.789% |
| Monocyte_LPS+DEX | 260756 | 7410 | 2.842% |
| Monocyte_PHA | 260756 | 26317 | 10.093% |
| Monocyte_PHA+DEX | 260756 | 8444 | 3.238% |
| NKcell_LPS | 259614 | 16 | 0.006% |
| NKcell_LPS+DEX | 259614 | 7297 | 2.811% |
| NKcell_PHA | 259614 | 1064 | 0.410% |
| NKcell_PHA+DEX | 259614 | 8788 | 3.385% |
| Tcell_LPS | 260818 | 337 | 0.129% |
| Tcell_LPS+DEX | 260818 | 11020 | 4.225% |
| Tcell_PHA | 260818 | 6633 | 2.543% |
| Tcell_PHA+DEX | 260818 | 17475 | 6.700% |
| DC_LPS | 211019 | 58 | 0.027% |
| DC_LPS+DEX | 211019 | 60 | 0.028% |
| DC_PHA | 211019 | 26 | 0.012% |
| DC_PHA+DEX | 211019 | 70 | 0.033% |

**Table S6:** Comparison of DARs increasing or decreasing chromatin accessibility for each condition. Column 2, number of DARs with decreased changes in treatment compared to the contrast group; Column 3, number of DARs with increased changes in treatment compared to the contrast group; Column 4 and 5 represent Fold changes between increase and decrease in chromatin accessibility ; and Column 6, p-value from single side binomial test.

| conditions | #DARs(LFC<0) | #DARs(LFC>0) | Fold decrease | Fold Increase | p-value |
| --- | --- | --- | --- | --- | --- |
| Bcell_LPS | 8 | 10 | 0.8 | 1.25 | 4.073e-01 |
| Bcell_LPS+DEX | 929 | 1395 | 0.67 | 1.5 | 1.869e-22 |
| Bcell_PHA | 679 | 1684 | 0.4 | 2.48 | 4.399e-98 |
| Bcell_PHA+DEX | 3326 | 1674 | 1.99 | 0.5 | 3.99e-123 |
| DC_LPS | 32 | 26 | 1.23 | 0.81 | 2.559e-01 |
| DC_LPS+DEX | 3 | 57 | 0.05 | 19 | 3.127e-14 |
| DC_PHA | 21 | 5 | 4.2 | 0.24 | 1.247e-03 |
| DC_PHA+DEX | 0 | 70 | 0 | NA | 8.47e-22 |
| Monocyte_LPS | 5122 | 9974 | 0.51 | 1.95 | <2.2e-16 |
| Monocyte_LPS+DEX | 3077 | 4333 | 0.71 | 1.41 | 1.139e-48 |
| Monocyte_PHA | 11475 | 14842 | 0.77 | 1.29 | 3.471e-96 |
| Monocyte_PHA+DEX | 4350 | 4094 | 1.06 | 0.94 | 2.758e-03 |
| NKcell_LPS | 2 | 14 | 0.14 | 7 | 2.09e-03 |
| NKcell_LPS+DEX | 3895 | 3402 | 1.14 | 0.87 | 4.163e-09 |
| NKcell_PHA | 217 | 847 | 0.26 | 3.9 | 1.125e-88 |
| NKcell_PHA+DEX | 5684 | 3104 | 1.83 | 0.55 | 2.331e-169 |
| Tcell_LPS | 64 | 273 | 0.23 | 4.27 | 3.581e-32 |
| Tcell_LPS+DEX | 5541 | 5479 | 1.01 | 0.99 | 2.806e-01 |
| Tcell_PHA | 986 | 5647 | 0.17 | 5.73 | <2.2e-16 |
| Tcell_PHA+DEX | 11583 | 5892 | 1.97 | 0.51 | <2.2e-16 |

**Table S7:** Summary of eGenes identified by different approaches at the same threshold (FDR*<*0.1 for each condition in the scRNAseq data. Column 2-6 represents the number of eGenes identified by `FastQTL`, `FastQTL` with 1000 permutations, `mashr`, `TORUS` and `DAP-G`. The number in bracket means the number of eGenes overlapped with those in `DAP-G`

| conditions | #eGenes (FastQTL) | #egene (FastQTL permutation) | #eGenes (mashr) | #eGenes (TORUS) | #eGenes (DAP-G) |
| --- | --- | --- | --- | --- | --- |
| Bcell_CTRL | 231 (25) | 12 (12) | 2649 (20) | 18 (18) | 25 |
| Bcell_LPS | 310 (30) | 18 (15) | 2656 (26) | 25 (25) | 32 |
| Bcell_LPS+DEX | 153 (28) | 10 (10) | 2650 (27) | 22 (22) | 29 |
| Bcell_PHA | 170 (31) | 14 (14) | 2652 (26) | 26 (26) | 32 |
| Bcell_PHA+DEX | 120 (22) | 9 (9) | 2678 (22) | 22 (22) | 28 |
| Monocyte_CTRL | 106 (17) | 6 (6) | 2603 (12) | 16 (15) | 19 |
| Monocyte_LPS | 161 (22) | 15 (13) | 2559 (17) | 21 (21) | 23 |
| Monocyte_LPS+DEX | 153 (27) | 15 (14) | 2586 (23) | 27 (25) | 32 |
| Monocyte_PHA | 141 (16) | 9 (8) | 2547 (15) | 17 (14) | 17 |
| Monocyte_PHA+DEX | 172 (25) | 15 (14) | 2590 (20) | 23 (23) | 29 |
| NKcell_CTRL | 212 (44) | 32 (25) | 2624 (44) | 35 (35) | 48 |
| NKcell_LPS | 213 (51) | 28 (26) | 2592 (51) | 44 (44) | 58 |
| NKcell_LPS+DEX | 38 (12) | 7 (7) | 2462 (15) | 13 (12) | 17 |
| NKcell_PHA | 53 (16) | 14 (13) | 2602 (21) | 17 (17) | 23 |
| NKcell_PHA+DEX | 59 (15) | 7 (7) | 2477 (15) | 13 (13) | 17 |
| Tcell_CTRL | 955 (248) | 186 (161) | 2605 (242) | 204 (203) | 269 |
| Tcell_LPS | 1032 (251) | 179 (159) | 2642 (238) | 206 (205) | 267 |
| Tcell_LPS+DEX | 576 (163) | 100 (92) | 2684 (157) | 134 (134) | 180 |
| Tcell_PHA | 592 (148) | 90 (79) | 2666 (149) | 123 (120) | 164 |
| Tcell_PHA+DEX | 635 (158) | 109 (95) | 2712 (153) | 124 (122) | 176 |
| total | 4081 (499) | 343 (268) | 2984 (423) | 385 (377) | 519 |

**Table S8:**
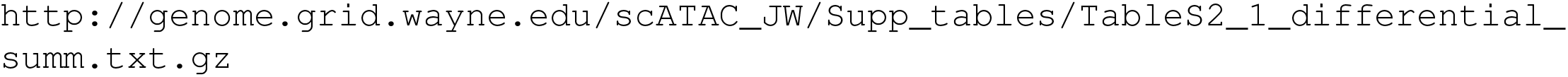
Results of differential chromatin accessibility analysis using DESeq2 in pseudo bulk aggregated data. Columns are: 1) Contrast; 2) Cell type; 3) Peak name; 4) baseMean, the average of the normalized count values across samples; 5) log_2_ fold change of chromatin accessibility for the peak; 6) Standard error; 7) *Z* − score 8) *p*-value; 9) FDR; and 10) combination of cell types and contrasts. http://genome.grid.wayne.edu/scATAC_JW/Supp_tables/TableS2_1_differential_summ.txt.gz

**Table S9:**
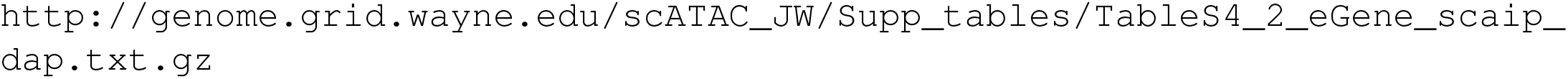
Fine-mapped eGenes identified by DAP-G at 10% FDR in single cell datasets. Columns are: 1) Ensembl gene ID; 2) local false discovery rate; 3) FDR; and 4) each dataset name, cell type and treatment. http://genome.grid.wayne.edu/scATAC_JW/Supp_tables/TableS4_2_eGene_scaip_dap.txt.gz

**Table S10:**
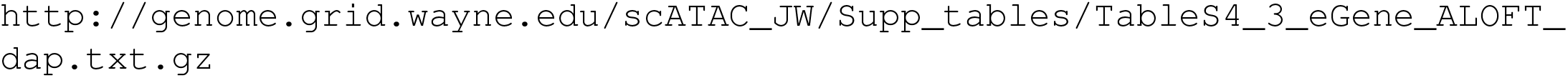
Fine-mapped eGenes identified by DAP-G at 10% FDR in the ALOFT whole blood dataset. Columns are: 1) Ensembl gene ID; 2) local false discovery rate; and 3) FDR. http://genome.grid.wayne.edu/scATAC_JW/Supp_tables/TableS4_3_eGene_ALOFT_dap.txt.gz

**Table S11:**
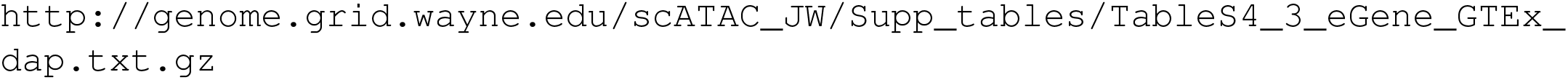
Fine-mapped eGenes identified by DAP-G at 10% FDR in the GTEx v8 whole blood dataset. Columns are: 1) Ensembl gene ID; 2) local false discovery rate; and 3) FDR. http://genome.grid.wayne.edu/scATAC_JW/Supp_tables/TableS4_3_eGene_GTEx_dap.txt.gz

**Table S12:** The results of protein coding genes associated with asthma risk with the combination of TWAS and colocalization analysis in ALOFT cohort. Columns are: 1) Ensembl gene ID; 2) Gene symbol; 3) The selected genetic variant for the SMR/TWAS analysis of the gene; 4) Chromosome; 5) chr pos grch38, the coordinate of genetic variant in hg38; 6) chr pos grch37, the coordinate of genetic variant in hg37 7) Posterior inclusion probability (PIP) of the SMR selected variant from textttDAP-G; 8) The *p*-values of eQTLs that are associated with gene expression from `fastQTL`; 9) genetic variants annotation; 10) TWAS *p*-values of gene associated with asthma risk ; 11) TWAS *Z*-score of gene associated with asthma risk; 12) FDR of TWAS analysis; 13) Gene locus-level colocalization probability (GLCP) from colocalization; 14) The probability of putative causal genes by combing TWAS and colocalization analysis using `INTACT`; 15) FDR from `INTACT` http://genome.grid.wayne.edu/scATAC_JW/Supp_tables/TableS5_1_asthma-risk-genes_ALOFT.txt.gz

**Table S13:** The results of protein coding genes associated with asthma risk with the combination of TWAS and colocalization analysis in GTEx whole blood tissue. Columns are: 1) Ensembl gene ID; 2) Gene symbol; 3) The selected genetic variant for the SMR/TWAS analysis of the gene; 4) Chromosome; 5) chr pos grch38, the coordinate of genetic variant in hg38; 6) chr pos grch37, the coordinate of genetic variant in hg37; 7) Posterior inclusion probability (PIP) of the SMR selected variant from textttDAP-G; 8) The *p*-values of eQTLs that are associated with gene expression from `fastQTL`; 9) Genetic variants annotation; 10) TWAS *p*-values of gene associated with asthma risk TWAS that are from the *p*-values of GWAS for the SMR selected variant; 11) TWAS *Z*-score of gene associated with asthma risk; 12) FDR of TWAS analysis; 13) Gene locus-level colocalization probability (GLCP) from colocalization; 14) The probability of putative causal genes by combing TWAS and colocalization analysis using `INTACT`; 15) FDR from `INTACT` http://genome.grid.wayne.edu/scATAC_JW/Supp_tables/TableS5_2_asthma-risk-genes_GTEx.txt.gz

**Table S14:** The 98 asthma risk genes identified by INTACT in ALOFT cohort at 10% FDR and also with the causal variants annotated in Response motifs. Columns are: 1) Ensembl gene ID; 2) Gene symbol; 3) The selected genetic variant for the SMR/TWAS analysis of the gene; 4) Chromosome; 5) chr pos grch38, the coordinate of genetic variant in hg38; 6) chr pos grch37, the coordinate of genetic variant in hg37; 7) Posterior inclusion probability (PIP) of the SMR selected variant from `DAP-G`; 8) The *p*-values of eQTLs that are associated with gene expression from `fastQTL`; 9) Genetic variants annotation; 10) TWAS *p*-values of gene associated with asthma risk TWAS that are from the *p*-values of GWAS for the SMR selected variant; 11) TWAS *Z*-score of gene associated with asthma risk; 12) FDR of TWAS analysis; 13) Gene locus-level colocalization probability (GLCP) from colocalization; 14) The probability of putative causal genes by combing TWAS and colocalization analysis using `INTACT`; 15) FDR from `INTACT`; 16) Chromatin regions covering the selected genetic variants; 17) The binding response motifs, including the binding score for reference and alternative allele; 18-20) the allelic-specific analysis information, including the detection sample, coefficient and *p*-values http://genome.grid.wayne.edu/scATAC_JW/Supp_tables/TableS5_3_asthma-risk-genes_in_response_ALOFT.txt.gz

**Table S15:** The 127 asthma risk genes identified by INTACT in GTEx whole blood tissue at 10% FDR and also with the causal variants annotated in Response motifs. Columns are: 1) Ensembl gene ID; 2) Gene symbol; 3) The selected genetic variant for the SMR/TWAS analysis of the gene; 4) Chromosome; 5) chr pos grch38, the coordinate of genetic variant in hg38; 6) chr pos grch37, the coordinate of genetic variant in hg37; 7) Posterior inclusion probability (PIP) of the SMR selected variant from `DAP-G`; 8) The *p*-values of eQTLs that are associated with gene expression from `fastQTL`; 9) Genetic variants annotation; 10) TWAS *p*-values of gene associated with asthma risk TWAS that are from the *p*-values of GWAS for the SMR selected variant; 11) TWAS *Z* − score of gene associated with asthma risk; 12) FDR of TWAS analysis; 13) Gene locus-level colocalization probability (GLCP) from colocalization; 14) The probability of putative causal genes by combing TWAS and colocalization analysis using `INTACT`; 15) FDR from `INTACT`; 16) Chromatin regions covering the selected genetic variants; 17) The binding response motifs, including the binding score for reference and alternative allele; 18-20) the allelic-specific analysis information, including the detection sample, coefficient and *p*-values http://genome.grid.wayne.edu/scATAC_JW/Supp_tables/TableS5_4_asthma-risk-genes_in_response_GTEx.txt.gz

